# *Multiple-Basin Gō-Martini* for Investigating Conformational Transitions and Environmental Interactions of Proteins

**DOI:** 10.1101/2024.12.11.628061

**Authors:** Song Yang, Chen Song

## Abstract

Proteins are inherently dynamic molecules, and their conformational transitions among various states are essential for numerous biological processes, which are often modulated by their interactions with surrounding environments. Although molecular dynamics (MD) simulations are widely used to investigate these transitions, all-atom (AA) methods are often limited by short timescales and high computational costs, and coarse-grained (CG) implicit-solvent Gō-like models are usually incapable of studying the interactions between proteins and their environments. Here, we present an approach called Multiple-basin Gō-Martini, which combines the recent Gō-Martini model with an exponential mixing scheme to facilitate the simulation of spontaneous protein conformational transitions in explicit environments. We demonstrate the versatility of our method through five diverse case studies: GlnBP, Arc, Hinge, SemiSWEET, and TRAAK, representing ligand-binding proteins, fold-switching proteins, *de novo* designed proteins, transporters, and mechanosensitive ion channels, respectively. The Multiple-basin Gō-Martini offers a new computational tool for investigating protein conformational transitions, identifying key intermediate states, and elucidating essential interactions between proteins and their environments, particularly protein-membrane interactions. In addition, this approach can efficiently generate thermodynamically meaningful datasets of protein conformational space, which may enhance deep learning-based models for predicting protein conformation distributions.

## Introduction

Large-scale conformational transitions of proteins play an essential role in living systems. Many proteins change their structure to perform a variety of functions. For example, ligand-binding proteins undergo conformational transitions from an open state to a closed state after capturing the substrate in the binding pocket. ^1,2^ Transporters alternatively open or enclose the outward and inward pockets to translocate the substrates across the membrane. ^3^ Ion channels can open the central pore and allow ions to pass through the membrane after being activated by various stimuli, such as membrane potential or ligand binding. ^4,5^ Thus, understanding the molecular mechanisms through conformational changes in proteins, especially membrane proteins, is becoming a fundamental and essential task in the current fields of structural biology and biophysics.

Molecular dynamics (MD) simulation is one of the most widely used computational methods, which bridges the gap between various experimental results and theoretical mechanisms underlying the intricate conformational changes of proteins. ^6–8^ The commonly used timescale for conventional all-atom MD simulations is around microseconds, while the conformational transitions of proteins in reality are usually in milliseconds or greater. ^9^ Therefore, although allatom MD simulations have achieved excellent success in describing protein dynamics, it is still challenging to sample large-scale conformational transitions of proteins, even with the use of MD-specialized supercomputers. ^10^ And the expensive computation cost also impedes the highthroughput application of MD simulations. Structure-based coarse-grained (CG) models are expected to overcome the above limitations by uniting groups of atoms into pseudoatoms and thus substantially reducing the computational complexity and enhancing the simulation speed. One of the most widely used CG models is the Gō-like model, such as the AICG2+ model ^11^ and the Karanicolas-Brooks Gō-like model, ^12,13^ which utilizes the Lennard-Jones-like potential network based on the contact map to maintain the second and higher structures and have achieved great success in simulating conformational changes of proteins using the switching method ^14–17^ and the multiple-basin approach. ^18–20^ However, the lack of an explicit solvent environment and biomembrane in these models may hinder the system from accurately simulating the conformational transition process of proteins, especially when the interactions between proteins and lipids are crucial, although much effort is being made in these directions. ^21–23^

Martini force field is one of the most popular CG force fields in simulating protein-lipid interactions, which has been systemically parameterized based on the partitioning free energies of a large number of chemical compounds between the water and oil phases. ^24,25^ However, classic Martini protein models need to use the elastic network potential to maintain protein stability, which inadvertently restricts protein conformational changes and limits the broader applications of the Martini force field. ^26^ Despite various attempts to overcome these limitations, ^27,28^ progress has been hindered by the inherent shortcomings of elastic network models in accurately representing protein conformational transitions.

Recently, the Gō-Martini model has emerged, which combines the Gō-like model with the Martini force field and can simulate large-scale dynamic transitions of proteins, resembling the traditional Gō-like models while preserving the advantages of the Martini force field in simulating protein-lipid interaction with an explicit environment. ^29–31^ Based on these promising characteristics of the Gō-Martini model, we have developed the Switching Gō-Martini approach to apply the switching method on the Gō-Martini model to achieve large-scale conformational transitions of proteins, especially membrane proteins (Fig. 1). ^32^ However, there are still some limitations in this approach, including the strong bias energy during the switching process and the nonspontaneous conformational transitions. ^14,32^ This hinders the study of the thermodynamics and kinetics of protein conformational transitions. Additionally, the secondary structure constraint in the Martini protein model hinders proteins from changing their conformations involving β-sheet structures. Therefore, in this work, we develop a method named Multiple-basin Gō-Martini, to overcome the above limitations by combining the multiple-basin approach and the Gō-Martini model.

**Figure 1:**
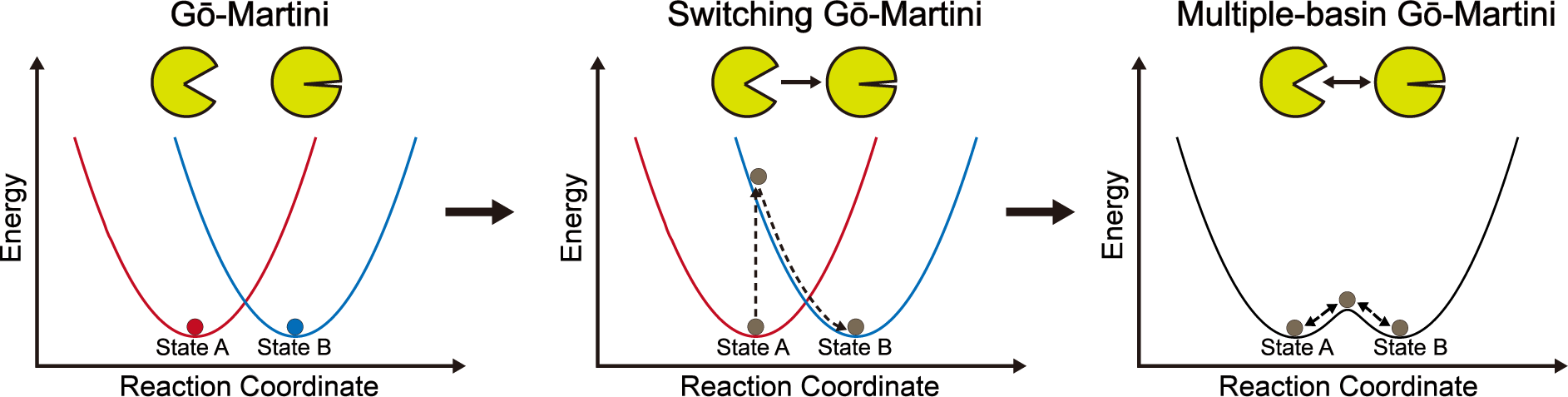
Development of Gō-Martini models for simulating protein conformational transitions. Energy landscapes are represented as simplified quadratic curves (blue and red) for Gō-Martini and Switching Gō-Martini models, or as black double-basin curves for Multiple-basin Gō-Martini model. Protein states are depicted by balls in red, blue, or brown. Yellow cartoons represent proteins, with arrows indicating the direction of conformational changes.

There are two primary approaches to combine single-basin potentials generated by structurebased Gō-like models to construct multiple-basin energy landscapes: the microscopic approach and the macroscopic approach. In the microscopic approach, each energy term of the multiplebasin potential is generated by combining the corresponding energy terms of individual singlebasin potentials. ^33–36^ This method effectively mixes potentials at the level of individual energy terms. In contrast, the macroscopic approach constructs the multiple-basin energy by using special functions to combine entire single-basin potentials, thus mixing at a macroscopic level. Currently, the macroscopic mixing scheme is more popular because of its requirement for fewer tunable parameters compared to the microscopic approach. Among macroscopic mixing functions, the exponential mixing scheme^18,20,37^ and the Hamiltonian mixing scheme^19,38,39^ are widely employed in multiple-basin simulations. For this study, we chose the exponential mixing scheme due to its simplicity and ease in defining two or more basins in the energy landscape. ^20^

Here we develop the łMultiple-basin Gō-Martiniž method by using the exponential macromixing scheme to investigate the conformational transitions of proteins, with a strong focus on membrane proteins. Firstly, We introduce the Multiple-basin Gō-Martini method and compare it with both the preliminary Gō-Martini model ^29,31^ and the Switching Gō-Martini method. ^32^ Then, we demonstrate the capability of our method using five case studies: (i) the closed–open transitions of glutamine binding protein (GlnBP), (ii) the sheet–helix fold-switching process of Arc, (iii) the unbinding and binding processes of the *de novo* designed protein Hinge, (iv) the conformational transition circle of the semiSWEET transporter, and (v) the surface tension-induced activation process of the mechanosensitive ion channel TRAAK. These results demonstrate that, with the Multiple-Basin Gō-Martini method, one can not only efficiently sample the conformational transition paths of proteins and identify the important intermediate states, but also gain semiquantitative insights into the dynamics, thermodynamics, and kinetics of protein conformations, as well as protein-protein and protein-membrane interactions.

## Results

### Multiple-basin Gō-Martini

The Multiple-basin Gō-Martini method employs an exponential mixing scheme to combine multiple single-basin potentials that represent different conformational states of a protein. This approach can construct a more complex and realistic energy landscape than Switching Gō-Martini, enabling proteins to oscillate between different potential basins and undergo conformational transitions within practicable simulation time. Fig. 1 illustrates the development of Gō-Martini models and their capabilities in simulating protein conformational transitions. In the classic Gō-Martini model, proteins are confined to a single basin of the energy landscape, defined by Lennard-Jones contact interactions. This restraint severely limits conformational transitions between multiple stable states. We recently proposed the Switching Gō-Martini method, which utilizes a simple and efficient approach to manually switch between energy landscapes of two protein states, enabling efficient simulations of conformational transitions. The Multiple-basin Gō-Martini method represents a significant advancement over these previous models, as proteins in the multiple-basin potential can spontaneously transition between energy basins without manual intervention, more closely mimicking natural protein dynamics. The exponential mixing function incorporates three key parameters - *β*, *C*_1_ and *C*_2_ - which can be fine-tuned to modulate the height of energy barriers and the relative depths of energy basins. More detailed implementations of the Multiple-basin Gō-Martini method are provided in the Materials and Methods section. To evaluate the sampling efficiency and reliability of our method, we have applied it to study five diverse systems, including GlnBP, Arc, Hinge, SemiSWEET, and TRAAK.

### Open–closed Transitions of GlnBP

GlnBP, a periplasmic binding protein that facilitates the uptake of L-glutamine in bacteria, is commonly used as a benchmark system to investigate protein conformational transitions. ^1^ GlnBP comprises two domains, a large domain and a small domain, connected by a hinge region. The crystal structures of GlnBP have been resolved both for the ligand-free open state and the ligand-bound closed state (Fig. 2a). ^40,41^ Many wet experiments and computational simulations have illustrated details of the open–closed transitions of GlnBP, making it an ideal benchmark system for validating the efficacy and efficiency of the Multiple-basin Gō-Martini method. ^42,43^

**Figure 2:**
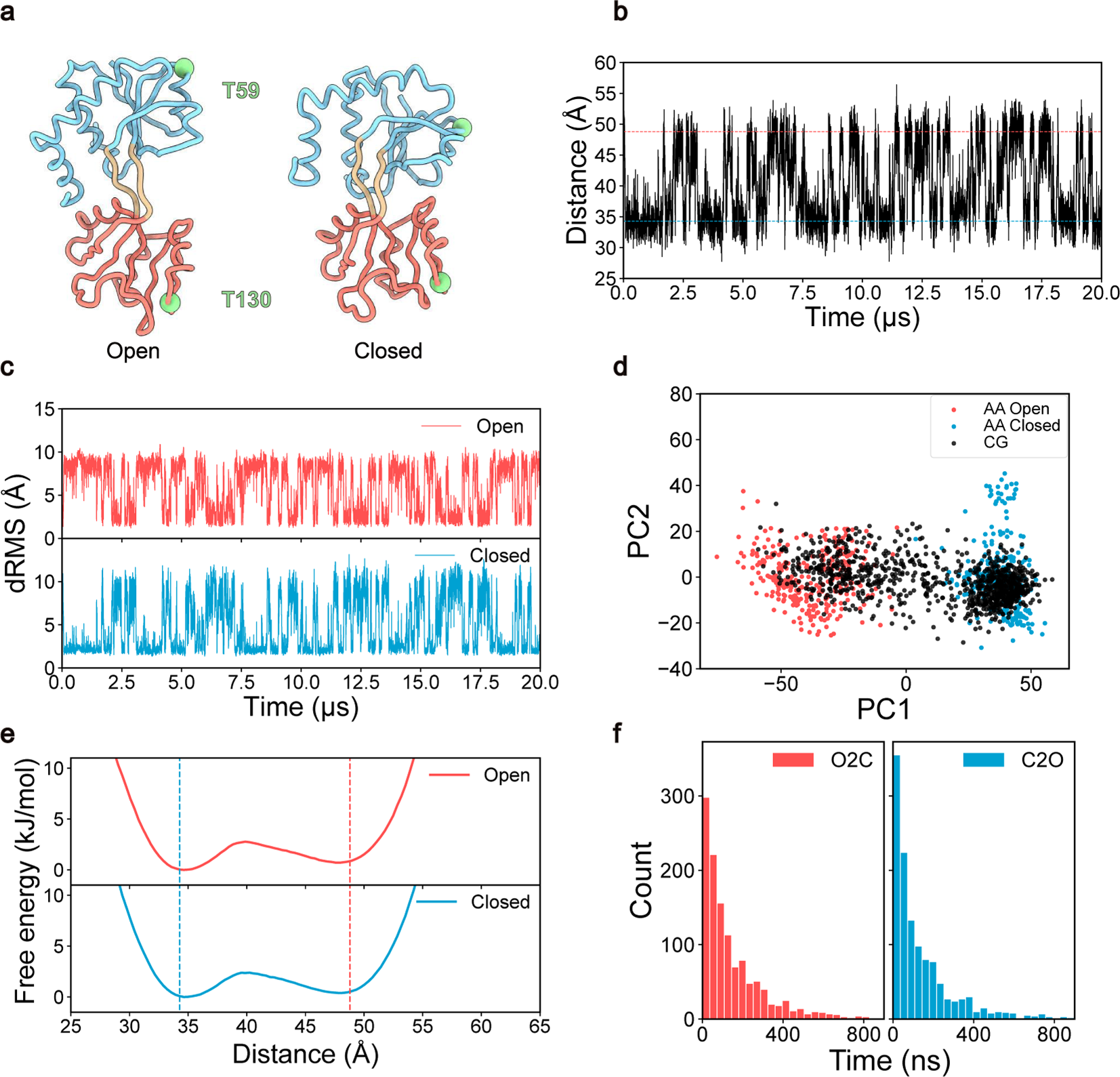
Multiple-basin Gō-Martini simulations for GlnBP conformational transitions. **a,** Coarse-grained structures of GlnBP in the open (PDB ID: 1GGG ^41^) and closed (PDB ID: 1WDN ^40^) states. The protein is shown as a ribbon, with the large domain colored sky blue and the small domain salmon. The remaining part is colored tan. The backbone atoms of T59 and T130 are highlighted with light green spheres. **b,** Time evolution of the distance between backbone atoms of T59 and T130 during one Multiple-basin Gō-Martini simulation. Red and blue dashed lines indicate reference distances from open and closed states, respectively. **c,** dRMS evolutions of GlnBP during the simulation, with respect to the open- (red) and closed- (blue) state structures. **d,** Principal component analysis (PCA) of GlnBP conformations. Coarse-grained simulation trajectories are projected onto the PC1-PC2 surface derived from all-atom simulations of open and closed GlnBP states. **e,** Free energy profiles of GlnBP from all CG simulations initiated from open and closed states, respectively. The first 1000 ns of each trajectory is discarded during the analysis. Red and blue dashed lines represent reference distances from open and closed state structures, respectively. The shaded area surrounding each line represents the statistical uncertainty of the free energy, estimated using bootstrap analysis. Note that the error is negligibly small and thus not visible in the figure. **f,** Distribution of transition times for open-to-closed (O2C) and closed-to-open (C2O) conformational transitions of GlnBP.

We constructed a multiple-basin Gō-Martini model for GlnBP, designating the open form as basin A and the closed state as basin B. To modulate the relative populations of open and closed states in our simulations, we fine-tuned the parameters *β*, *C*_1_ and *C*_2_ based on the experimental state distribution values of GlnBP. In this study, we set all *C*_2_ values to 0 and systematically adjusted *β* and *C*_1_. T59 and T130 of GlnBP are commonly used as fluorescence labeling sites in single-molecule Förster resonance energy transfer (smFRET) experiments. ^42,43^ Therefore, we utilized the distance between the backbone (BB) atoms of T59 and T130 as the reaction coordinate to characterize the conformational changes of GlnBP during our parameter search. Various parameter combinations were systematically explored to identify the optimal set for our MD simulations of GlnBP. We investigated five values for parameter *β* (1/100, 1/200, 1/300, 1/400, and 1/500 mol/kJ) and five values for parameter *C*_1_ (-100, -200, -300, -400, and -500 kJ/mol), resulting in a total of 25 different parameter combinations. For each combination, a 5-µs simulation was conducted. The results of these simulations are presented in Supplementary Fig. S1. Our analysis revealed that the parameter *C*_1_ predominantly influenced the relative population of conformational states. More negative *C*_1_ values were correlated with an increased prevalence of open conformations, while more positive *C*_1_ values favored closed states. The parameter *β*, on the other hand, modulated the energy barrier between conformational states. As *β* varied from 1/100 to 1/500 mol/kJ, we observed the emergence of conformational transitions and intermediate states in our simulations. Interestingly, *β* not only affected the barrier in the energy landscape of GlnBP, but also influenced the relative populations of open and closed states (Supplementary Fig. S1). This suggests a complex interplay between the two parameters in determining the conformational landscape of GlnBP.

Following a comprehensive parameter search, we identified the optimal parameter combination (*β* =1/300 mol/kJ, *C*_1_=-300 kJ/mol, *C*_2_=0 kJ/mol) for the Multiple-basin Gō-Martini simulations of GlnBP. Using this set of parameters, we conducted a 20-µs simulation, during which we observed reversible transitions between different conformational states of GlnBP (Supplementary Movie S1). As shown in Fig. 2b, the distance changes between residues T59 and T130 reveal repeated cycles of pocket opening and closing throughout the simulation. To characterize the overall conformational changes of GlnBP, we employed distance root-mean-square displacement (dRMS) as a reaction coordinate, which calculated the changes of residue-residue distance along the simulations with the difference between two states over 5 Å. This metric provides a representation of structural changes across the entire protein. As illustrated in Fig. 2c, the dRMS analysis clearly demonstrates that GlnBP underwent spontaneous and dynamic conformational transitions between open and closed states with our Multiple-basin Gō-Martini method and without the need for external manipulations as in the Switching Gō-Martini method.

Principal component analysis (PCA) is one of the most widely used dimension reduction methods, capable of extracting the principal motions from simulation trajectories. ^44^ In this study, we applied PCA to analyze all-atom MD simulations of the open and closed states of GlnBP, projecting the CG simulation results onto these PCA modes to assess the similarity between coarse-grained and all-atom simulations. We conducted three independent all-atom simulations for both the open and closed states of GlnBP, with each trajectory spanning 1000 ns. In our analysis, we focused solely on the center of mass of the backbone of each residue in the all-atom simulations, in accordance with the projection rules of the Martini coarse-graining strategy. The results presented in Fig. 2d indicate that the first (PC1) and second (PC2) principal components from the all-atom simulations closely align with those from the coarse-grained simulations. PC1 corresponds to the open and closed motions of the GlnBP binding pockets, accounting for 62.1% of the total variance. Meanwhile, PC2 reflects the relative twisting motions of the large and small domains of GlnBP, occupying 7.3% of the variance. Overall, these findings demonstrate that our Multiple-basin Gō-Martini simulations can effectively capture the key motions observed in the all-atom MD simulations.

To assess the impact of initial conformations on the equilibrium distribution of states, we conducted two sets of simulations using the open and closed states of GlnBP as starting points, respectively. For each initial state, we performed 10 independent simulations, each running for 20 µs, resulting in a total of 200 µs per initial condition. The Supplementary Fig. S2 displays the time series of the binding pocket distance for each trajectory, in which the diversity between each trajectory confirms the independence of each simulation. We aggregated data from all trajectories to compute the free energy profiles for each initial state condition. Our analysis reveals that simulations initiated from the open state yield a conformational equilibrium constant (*K_Closed/Open_*) of 0.89, while those starting from the closed state produce a value of 0.83 (Fig. 2e). The similarity between these equilibrium constants suggests that the choice of the initial conformation does not significantly bias the long-term equilibrium distribution of states.

The conformational equilibrium constant derived from all the 20 simulation trajectories was 0.84, falling within the range observed in the smFRET experiments (0.64-1.01). ^42,45^ Having established the model’s thermodynamic accuracy, we proceeded to evaluate its kinetic properties. Using the rate constants obtained from Markov MD simulations of GlnBP, ^43^ we estimated the average transition time from closed to open states to be 14.5 µs, while the transition time from open to closed states was found to be 1.0 µs in the presence of the ligand. To estimate the conformational transition time in our simulations, we employed a method in which open-to-closed transitions begin when the distance between T59 and T130 leaves the open basin value and end upon reaching the closed basin value. ^6^ The closed-to-open transitions follow the reverse process. Fig. 2f illustrates the distribution of the calculated transition times, in which we observed 1225 closed-to-open transitions and 1228 open-to-closed transitions in our total 400 µs simulations. Using the above method, we calculated the average open-to-closed transition time as 157 ± 186 ns, and the closed-to-open transition time as 149 ± 191 ns. Notably, the transition rates in our simulations are 6–97 times faster than those observed in all-atom simulations. This acceleration is consistent with expectations for coarse-grained simulations, which simplify the protein energy landscape. However, it is important to note that our coarse-grained simulations do not account for ligand binding effects. Additionally, our Multiple-basin Gō-Martini method intentionally underestimates energy barriers to facilitate conformational changes within the given simulation timescales. Consequently, this approach enables the observation of GlnBP conformational dynamics, albeit at an accelerated pace compared to fully detailed all-atom simulations.

### Fold-switching Process of Arc

The wild-type Arc repressor is a homodimer characterized by a β-sheet structure at its N-terminus. ^46^ Interestingly, a double mutant of the Arc repressor containing N11L and L12N substitutions is reported to adopt a novel native fold, in which the N-terminal structure transits from an antiparallel β-sheet to two 3_10_ helices. ^47^ Additionally, the intermediate single mutant N11L was found to populate both the sheet and helix structures (Fig. 3a), exhibiting the ability to interchange between these conformations. ^18,48^ This conformational flexibility of the N11L mutant presents an excellent model system to examine the ability of the Multiple-basin Gō-Martini method in simulating transitions between protein secondary structures.

**Figure 3:**
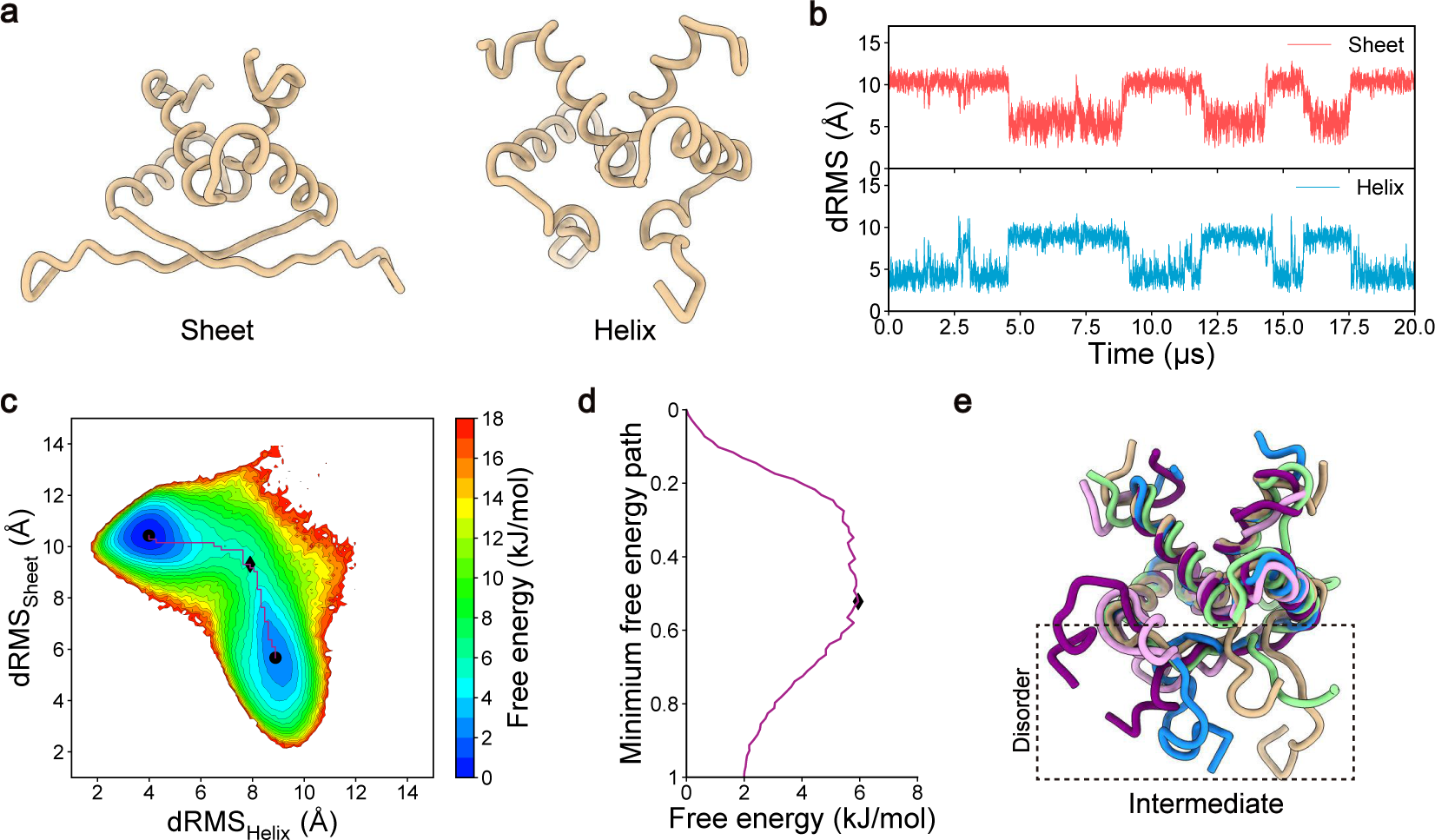
Multiple-basin Gō-Martini simulations for Arc conformational transitions. **a,** Coarsegrained structures of Arc in the sheet (PDB ID: 1ARR ^46^) and helix (PDB ID: 1QTG ^47^) states. The protein is shown as a tan ribbon. **b,** dRMS evolutions of Arc during the simulation, relative to the sheet (red) and helix (blue) reference structures. **c,** Free energy surface of Arc plotted as a function of dRMS values with respect to the sheet and helix states. Data from all trajectories, excluding the initial 1000 ns of each, are used for analysis. Black circles and diamond represent the steady basin states and intermediate states, respectively. The purple curve delineates the minimum free energy path. **d,** Free energy profile along the minimum free energy path. The black diamond represents the intermediate states. **e,** Representative intermediate structures of Arc at the transition barrier. Five distinct intermediate structures are illustrated here, each represented in a different color.

To investigate this system, we used Gō-Martini models to generate single-basin potentials for the N11L Arc mutant in both N-terminal β-sheet and helix conformations. We then utilized the exponential macro-mixing method to construct a double-basin energy landscape by combining these two single basins. Following a parameter optimization process similar to that used for GlnBP, we identified a set of optimized parameters that successfully reproduced spontaneous transitions between the β-sheet and 3_10_ helix conformations (Supplementary Movie 2). With the helix structure as the initial state, we conducted 10 independent simulations, each running for 20 µs, resulting in a total simulation time of 200 µs. Using the dRMS corresponding to the sheet and helix states as the reaction coordinates to characterize the conformational changes of Arc, we observed clear oscillations, indicating that Arc indeed underwent spontaneous conformational transitions between the β-sheet and helix states (Fig. 3b, Supplementary Fig. S3).

To characterize the state transitions of the mutant Arc, we utilized a two-dimensional (2D) free energy surface to visualize the conformational changes. As shown in Fig. 3c, two distinct energy basins are evident. The first basin is located at dRMS_Helix_ = 3.99 Å and dRMS_Sheet_ = 10.43 Å, while the second basin is located at dRMS_Helix_ = 8.89 Å and dRMS_Sheet_ = 5.67 Å. The slightly larger dRMS for the stable sheet and helix conformations can be attributed to the flexibility of the Nand C-termini of the protein. We extracted the representative structures of the two basins and aligned them to the reference structures (Supplementary Fig. S4). When excluding the flexible terminal regions (residues 1-8 and 49-53), the root mean square deviation (RMSD) of them was 1.95 and 1.83 Å for the reference sheet and helix states, respectively. Based on the barrier positions identified through dRMS_Helix_ and dRMS_Sheet_, we roughly partitioned the free energy surface into four regions. Analysis of the relative populations between the two major basins (located in the right-bottom and left-top regions) yielded a conformational equilibrium constant (*K_Sheet/Helix_*) of 0.71. The N11L mutant of Arc has been reported to sample both the helix and the sheet states, with the helical dimer predominating at a ratio of approximately 2:1 or 3:1 over the sheet form at 25°C. ^48,49^ Our result closely aligns with these observations.

Upon further analysis of the free energy landscape, we identified a distinct transition pathway connecting two basins (Fig. 3c). By plotting the minimum free energy profile along this pathway, we observe that Arc underwent a conformational transition through an intermediate state distant from both the helix and the sheet states (Fig. 3d). The energy barrier peak is located at dRMS_Helix_ = 7.91 Å and dRMS_Sheet_ = 9.31 Å. As illustrated in Fig. 3e, the N-terminus of the intermediate state exhibits disordered structure, suggesting that Arc follows a helix-disorder-sheet transition. These results demonstrate the capability of the Multiple-basin Gō-Martini method to elucidate the conformational transition of fold-switching proteins.

### Substrate Unbinding and Binding of de novo-designed Hinge

*De novo* protein design has emerged as a frontier in protein engineering. It is intriguing to evaluate whether our Multiple-basin Gō-Martini method can be applied to *de novo* designed proteins. To address this, we investigated the conformational transitions of a recently designed two-state hinge protein, ^50^ cs207 (hereafter referred to as Hinge), with particular interest in its substrate peptide unbinding and binding dynamics. Hinge exhibits two distinct conformational states: a bound state, where the binding pocket is accessible to bind the substrate peptide, and an unbound state, where the binding pocket is enclosed and the peptide is displaced (Fig. 4a). To elucidate the molecular mechanisms underlying these transitions, we employed steered MD as well as unbiased simulations to characterize the unbinding and binding processes between Hinge and its substrate peptide.

**Figure 4:**
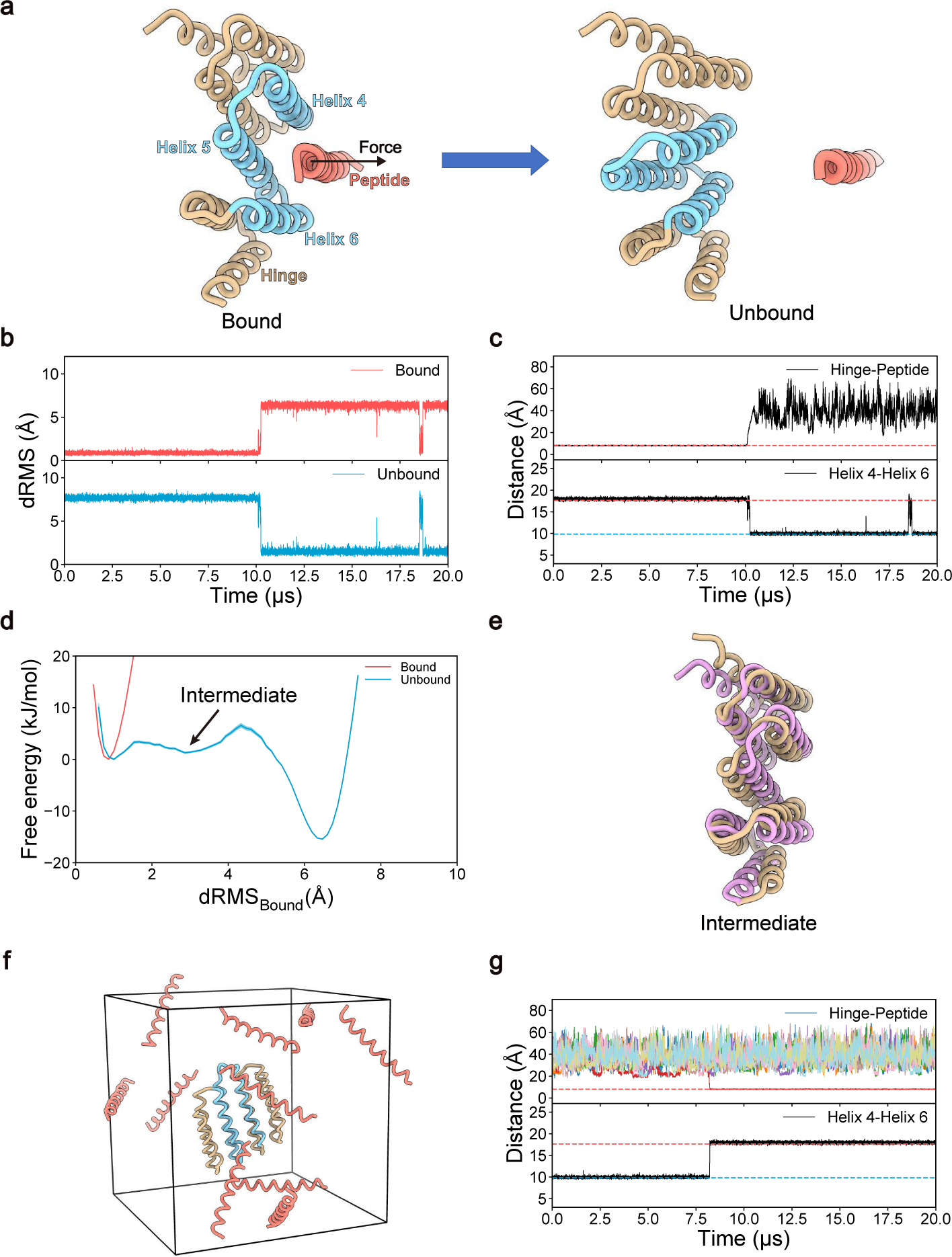
Multiple-basin Gō-Martini simulations for conformational transitions of *de novo* designed Hinge protein. **a,** Coarse-grained structures of Hinge in the bound (PDB ID: 8FIQ) and unbound (PDB ID: 8FIN) states. ^50^ The backbone of Hinge is shown in tan, with Helices 4-6 highlighted in sky blue. The substrate peptide is depicted as a salmon ribbon. **b,** dRMS evolutions of Hinge during the pulling simulations, relative to the bound (red) and unbound (blue) reference structures, respectively. **c,** Time evolution of the distance between Hinge and substrate peptide (upper panel) and the distance between Helix 4 and Helix 6 (lower panel). The dashed lines indicate the corresponding reference distance in the bound (red) and unbound (blue) states, respectively. **d,** Free energy profiles of Hinge conformations under peptide-bound and peptide-unbound conditions. Analysis of MD trajectories spanning 1–10 µs across 10 independent replicates reveals the conformational distribution of Hinge under the peptide-bound condition (red curve), while the subsequent 11–20 µs simulation data characterizes the free energy profile of Hinge under the peptide-unbound condition (blue curve). The shaded area surrounding each line represents the statistical uncertainty of the free energy, estimated using bootstrap analysis. **e,** Representative intermediate conformation of Hinge (tan ribbon) superimposed with the reference bound state (plum). **f,** Simulation system showing Hinge in the unbound state with ten randomly distributed substrate peptides around. **g,** Time evolution of Hinge-peptide distances (upper panel, individual traces for each peptide) and Helix 4-Helix 6 distance (lower panel). The dashed lines indicate the corresponding reference distances in the bound (red) and unbound (blue) states, respectively.

We constructed a dual-basin energy landscape of the Hinge protein using the exponential mixing approach, initializing the simulation system with the substrate peptide bound to Hinge. Initially, we performed 10 µs of standard MD simulations to characterize the bound-state dynamics of the Hinge-peptide complex. Subsequently, we employed steered MD with a constant-velocity force applied to the centers of mass of the Hinge binding pocket (Helices 4-6, residues 67-132) and its substrate peptide, separating them by 4 nm over 500 ns (Fig. 4a). Following this, we conducted standard MD simulations extending to 20 µs. This process was repeated in 10 independent simulations. As illustrated in Fig. 4b and Supplementary Fig. S5, the dRMS with respect to the bound and unbound states of Hinge demonstrates that the Hinge maintained the bound conformation when complexed with the peptide. Upon forced separation of the substrate peptide, Hinge underwent a a conformational transition from the bound to the unbound state (Supplementary Movie S3). We characterized the binding pocket conformation using the distance between Helix 4 and Helix 6 (Fig. 4c). This analysis shows that the binding pocket adopts an open configuration when the substrate peptide is bound and closes upon peptide release. Across all 10 independent simulations, the Hinge-peptide complex exhibited a single, well-defined bound conformational state. In contrast, during the unbound phase, while Hinge sampled both conformational states, the bound configuration was rarely observed, occupying less than 1% of the total simulation time.

Using dRMS_Bound_ as the reaction coordinate, free energy analysis demonstrates that peptide binding significantly modulates the conformational distribution of Hinge (Fig. 4d). In the absence of peptide binding, the unbound state of Hinge exhibits a -15.4 kJ/mol energy difference relative to the bound state. With substrate peptide bound, we did not observe spontaneous unbinding events, indicating that the binding affinity between Hinge and the peptide is very high, and peptide binding stabilizes Hinge in this single bound state. Our analysis of the conformational distribution of Hinge during the peptide-unbound condition identified an intermediate state. As shown in Fig. 4e, this intermediate state resembles the bound conformation of Hinge, but exhibits a partially closed binding pocket, where the distance between Helix 4 and Helix 6 is 13.6 Å, compared to 17.7 Å in the fully bound state.

In all 10 replicates of the unbinding simulations, we did not observe spontaneous re-binding events. To further investigate the binding mechanism, we constructed a second system comprising Hinge in its unbound state with 10 substrate peptides randomly distributed in the simulation box (Fig. 4f). We performed 10 independent simulations, each running 20 µs, to study the binding process. Among these simulations, we observed five successful binding events (Supplementary Fig. S5). Fig. 4g illustrates a representative binding trajectory, where one of the 10 peptides successfully docked into the binding pocket of Hinge when the pocket adopted an open conformation, subsequently stabilizing Hinge in its bound state (Supplementary Movie S4). These complementary unbinding and binding simulations demonstrate that the Multiple-basin Gō-Martini method can effectively capture the conformational dynamics of *de novo* designed proteins and elucidates the molecular mechanisms underlying the binding-unbinding transitions of protein-peptide and protein-protein complexes.

### Transport Cycle of SemiSWEET

The fourth case study focuses on the minimal sugar transporter SemiSWEET. Transporters play a crucial role in cellular function by facilitating the translocation of small molecules across biological membranes. This process is essential for the absorption of nutrients and the expulsion of potentially hazardous molecules. The widely accepted mechanism for substrate transport is the alternating access model. ^3,51^ In this model, the outward-facing and inward-facing binding pockets alternately open and close, allowing substrates to move across the membrane as these binding sites change conformations. This process typically involves three distinct conformations: outward-open, occluded, and inward-open states, where the outward and inward binding sites are either accessible or closed off. SemiSWEET represents one of the smallest known transporters, with a molecular mass less than 20 kDa and a structure comprising two symmetrical units. ^52–54^ It functions by transporting sugar molecules from the extracellular into the intracellular sides of a membrane along the concentration gradient. Importantly, the structures of both the outward-open and inward-open states of SemiSWEET have been experimentally resolved. ^53^ Given these structural insights, we aim to utilize SemiSWEET as a model system to evaluate the efficacy of our Multiple-basin Gō-Martini approach in simulating conformational transitions of membrane proteins.

We used the Multiple-basin Gō-Martini method to construct a double basin energy landscape with crystal structures of the outward-open and inward-open states as the reference(Fig. 5a). We performed coarse-grained MD simulations of SemiSWEET from the outward-open state (10 simulations totaling 200 us) after parameter optimization. In all simulations, we find that SemiSWEET can change its conformations and complete the translocation cycle within the limited 20 µs timescale (Supplementary Movie S5). In Fig. 5b, we can see the dRMS with respect to the outward-open and inward-open states of SimiSWEET changes along the simulation time, which clearly presents the conformational transition process between the two states. Because dRMS only reflects the overall changes of proteins and cannot reveal the local details, we analyzed the width of the outward and inward binding pockets (distances of D59-D59, M39-M39 between chain A and B, respectively). Fig. 5c and Supplementary Fig. S6 show that the outward and inward binding pockets indeed alternatively open or close during the simulations.

**Figure 5:**
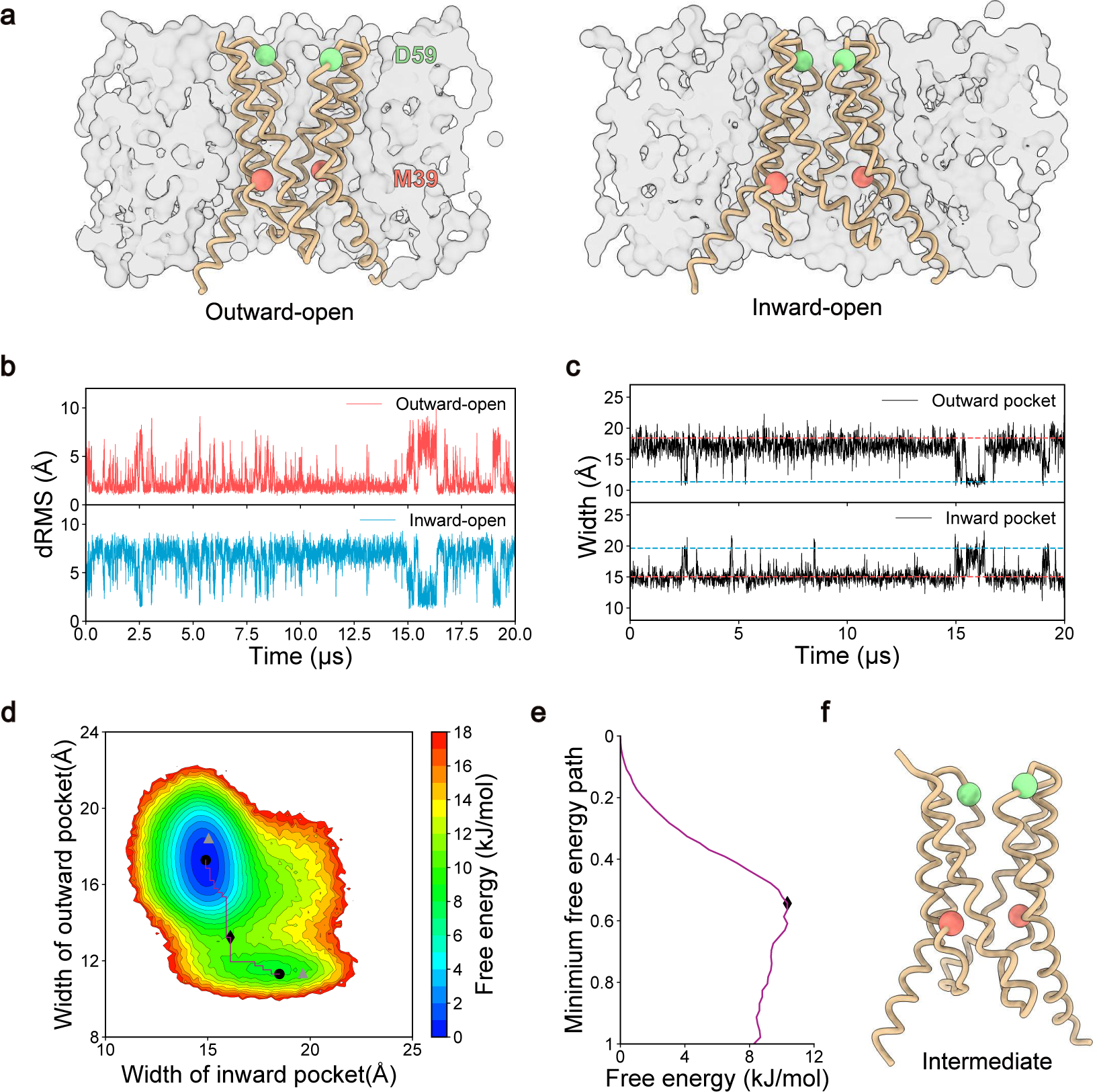
Multiple-basin Gō-Martini simulations for SemiSWEET conformational transitions. **a,** Coarse-grained structures of SemiSWEET in the outward-open (PDB ID: 4X5N) and inward-open (PDB ID: 4X5M) states ^53^ embedded in a lipid bilayer. The protein is shown as a tan ribbon, with the backbone atoms of M39 and D59 highlighted with salmon and light green spheres, respectively. The membrane is represented by gray surfaces. **b,** dRMS evolutions of SemiSWEET during the simulation, relative to the outward-open (red) and inward-open (blue) reference structures, respectively. **c,** Time evolution of the width of outward and inward binding pockets of SemiSWEET. The red and blue dashed line represent the width of corresponding binding pockets in the outward-open and inward-open states, respectively. **d,** Free energy landscape of SemiSWEET plotted as a function of width of outward and inward binding pockets as reaction coordinates. Data from all trajectories, excluding the initial 1000 ns of each, are used for analysis. Black circles and diamonds represent the steady basin states and intermediate states, respectively. The grey triangles represent the crystal structures. The purple curve delineates the minimum free energy path. **e,** Free energy profile along the minimum free energy path. The black diamond represents the intermediate states. **f,** Representative intermediate structure of SemiSWEET at the transition barrier.

To further elucidate the conformational changes of SemiSWEET, we constructed a 2D free energy surface using the widths of the outward and inward binding pockets as reaction coordinates. As illustrated in Fig. 5d, the free energy landscape reveals two distinct basins. The basin corresponding to the outward-open state is located at (14.9 Å, 17.3 Å), with the values representing the widths of the inward and outward binding pockets, respectively. In contrast, the basin denoting the inward-open state is positioned at (18.5 Å, 11.3 Å). These distances are closed to those of the reference structures, which show widths of (15.0 Å, 18.4 Å) for the outward-open state and (19.6 Å, 11.4 Å) for the inward-open state. Then we extracted the representative states from these two basins and compared them with the reference states (Supplementary Fig. S7). The RMSD between the computational and experimental structures was 1.94 Å for the outward-open state and 1.88 Å for the inward-open state. These results indicate that the basin structures we captured closely match the experimentally determined configurations. By mapping the minimum free energy path (Fig. 5d, e), we observe that the conformational transition follows a specific sequence from the outward-open to the inward-open states: initially, the outward binding pocket closes, followed by the opening of the inward binding pocket. This pathway suggests the presence of an intermediate occluded state, approximately located at (16.1 Å, 13.0 Å)(Fig. 5f). Notably, the alternating access mechanism observed in our simulations is consistent with previous experimental and simulation results, ^53,54^ which validates the efficacy and accuracy of our Multiple-basin Gō-Martini method in simulating membrane proteins.

### Mechanical Activation of TRAAK

Mechanosensitive ion channels play a crucial role in transducing external physical stimuli into cellular electrical signals by modulating their gating in response to mechanical forces. TRAAK, a member of the two-pore domain K^+^ (K2P) channel family, is one such mechanosensitive channel expressed in the nervous system. These channels contribute to the leak K^+^ conductance and help maintain the resting potential of neurons. ^55,56^ Under resting conditions, TRAAK exhibits low basal leak activity, mainly due to lipid occlusion of the pore. ^57^ However, when surface tension is applied to the membrane surrounding TRAAK, it induces conformational changes in the channel, which can enhance TRAAK activity up to ∼100-fold. ^57–59^ Specifically, movement of transmembrane helix (TM) 4 from a downward state to an upward state seals the fenestration of TRAAK, preventing lipid intrusion into the ion-conducting pore. ^57^ In addition, some work has recently pointed out that conformational changes in TM4 also modulate the states of the selective filter (SF) of K2P channels from low conductance to high conductance. ^60–62^ While MD simulations have been used to investigate the detailed process of how mechanical stimuli alter the states of K2P channels, ^60,63^ two significant questions remain: (1) The mechanical stimuli used in simulations are often exaggerated compared to physiological conditions (∼50 mN/m in simulations ^60^ vs. 0-10 mN/m in biological systems^64^). (2) A comprehensive depiction of the entire transition cycle of TRAAK and the influence of various surface tensions on state distribution using MD simulations is lacking. To address these gaps, we employ our newly developed Multiple-basin Gō-Martini method to reveal a more complete picture of TRAAK behavior under various surface tensions and elucidate the lipid interactions involved in this process.

We used the down and up conformational states of TRAAK as reference structures to construct a double-basin energy landscape using the Multiple-basin Gō-Martini approach (Fig. 6a). In the CG system, we embedded the membrane protein in a pure POPC bilayer. To simulate mechanical stimuli, we applied a surface tension of 10 mN/m to the membrane. After parameter optimization, we successfully simulated spontaneous conformational transitions of TRAAK (Supplementary Fig. S8 and Supplementary Movie S6). Fig. 6b illustrates the oscillations of the dRMS with respect to the up and down states over the courses of our 20-µs simulations. These oscillations clearly indicate that TRAAK underwent transitions between the two conformational states. To elucidate the critical motions of TM4, we selected the distance between the BB atoms of P155 (in TM2) and I279 (in TM4) as a reaction coordinate. As depicted in Fig. 6c, this distance oscillates between 6.8 Å and 11.9 Å, which aligns with the range of 6.6-12.4 Å observed in the reference structures for the up and down states. By analyzing the distance distribution between P155 and I279, we identified three distinct conformational states of TRAAK, representing down (11.9 Å), intermediate (8.3 Å), and up (6.8 Å) states, respectively(Fig. 6d). These results provide compelling evidence that our method can effectively simulate the up–down transitions of TRAAK under a membrane surface tension.

**Figure 6:**
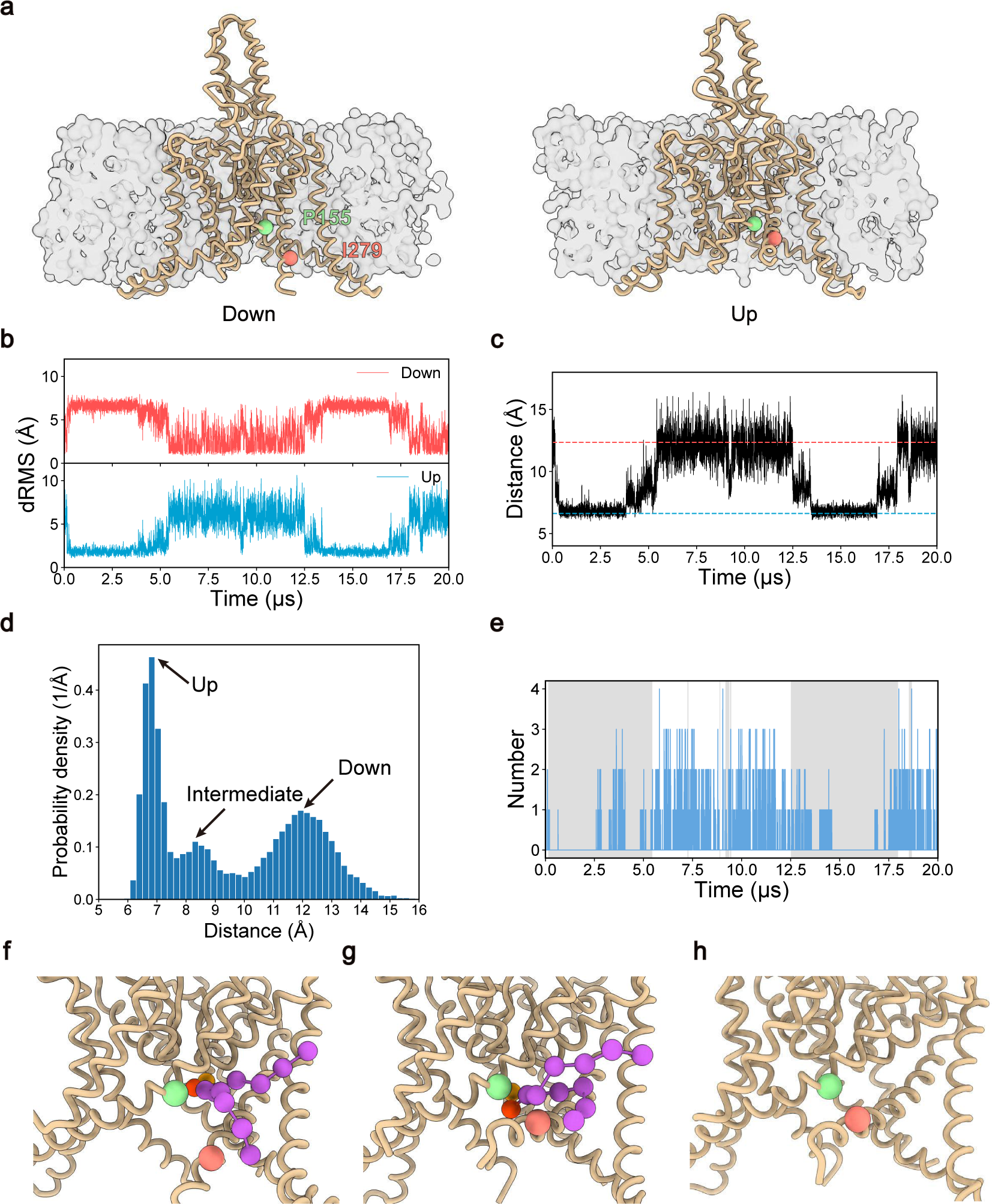
Multiple-basin Gō-Martini simulations for TRAAK conformational transitions. **a,** Coarse-grained structures of TRAAK in the down (PDB ID: 7LJB^65^) and up (PDB ID: 4WFE^57^) states embedded in a lipid bilayer. The protein is shown as a tan ribbon with the membrane represented by gray surfaces. The backbone atoms of P155 and I279 are highlighted with light green and salmon spheres, respectively. **b,** dRMS evolutions of TRAAK with a surface tension of 10 mN/m during the simulation, relative to the down (red) and up (blue) reference structures. **c,** Time evolution of distance between backbone atoms of P156 and I279 with a surface tension of 10 mN/m during a Multiple-basin Gō-Martini simulation. Red and blue dashed lines indicate reference distances from down and up states, respectively. **d,** Distribution of the P156-I279 distance during TRAAK conformational transitions. The histogram reveals three distinct conformational states representing up, intermediate, and down states, respectively. **e,** Time course of the number of lipids within an 8 Å radius from the center of the bottom region of the selective filter (T129 and T238 of chain A and B) in TRAAK with a surface tension of 10 mN/m during the simulation. The grey shaded area denotes the period when TRAAK is in the up state, while the white area the down state. **f-h,** Detailed views of TRAAK in the down (**f**), intermediate (**g**), and up (**h**) states with the presence or absence of bound lipids in the fenestration. The lipids are depicted using a ball-and-stick model, with the head groups colored orange and red while the remaining part colored middle orchid.

To investigate the interactions between lipids and TRAAK during conformational changes, we analyzed the number of lipid within an 8 Å radius from the center of the bottom region of the selective filter (T129 and T238 of chain A and B) in TRAAK. Our results reveal a significant decrease in the number of selected lipids when TRAAK transitioned to the up state, while an increase was observed in the down state (Fig. 6e). Traditionally, lipid tails are thought to be the primary cause to occlude the central ion-conducting pore of K2P channels, ^57^ while recent findings support a significant role for lipid heads. ^66,67^ To address the question of which lipid components insert into the fenestration of TRAAK and occlude the central pore, we examined the occupancy of lipid heads and tails in the central pore in both down and up states. Results from all 10 independent simulations show that lipid head group occupancy was 37 ± 7% in the down state and 9 ± 4% in the up state (Supplementary Fig. S9). In contrast, lipid tail occupancy remained negligible in both states. These findings clearly demonstrate that in the Martini force field, lipid heads rather than lipid tails are predominantly oriented toward the central pore of TRAAK by being inserted into the fenestration formed by TM2 and TM4. To elucidate the intricate interactions between lipids and different conformational states of TRAAK, we extracted representative structures of the down, intermediate, and up states based on the distance distribution between P155 and I279 (Fig. 6d). Fig. 6f-h illustrates representative zoom-in structures of TRAAK in these three states with or without lipids occupying the fenestration formed by TM2 and TM4. These images reveal that during the upward motion of TM4, lipid heads within the fenestration are displaced away from the center, which can facilitate ions passing through the channel. This lipid rearrangement associated with TRAAK conformational transitions appears to play a crucial role in modulating channel function.

To elucidate the influence of various surface tensions on the state distribution of TRAAK, we conducted a series of MD simulations with surface tensions ranging from 0 to 30 mN/m (0, 5, 10, 20, and 30 mN/m) (Fig. 7a). For each condition, we performed 10 independent 20-µs simulations. Fig. 7b displays the representative time series of dRMS with respect to the down state under distinct surface tensions. The results clearly demonstrate that with increasing surface tensions, the up state of TRAAK gradually becomes the predominant conformation during the simulations. To quantitatively illustrate the overall distributions of TRAAK conformations, Fig. 7c shows the free energy profiles along the dRMS with respect to the down state of TRAAK. Our results demonstrate that the free energy of the up state, with a dRMS value of about 6.8 Å, changes from approximately 3.6 to -4.9 kJ/mol as the surface tension increases from 0 to 30 mN/m. This indicates that the up state becomes favorable with increased membrane surface tension.

**Figure 7:**
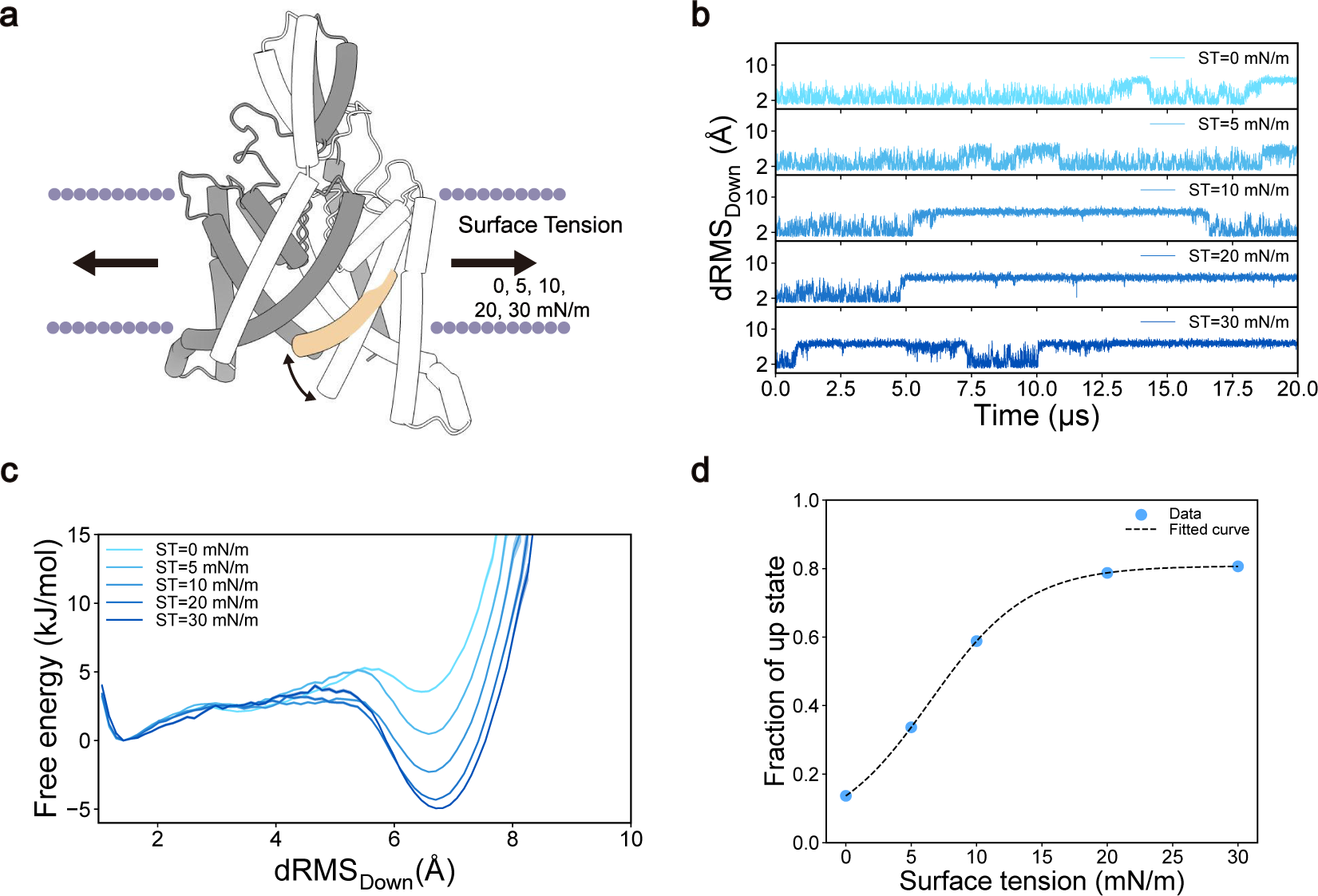
The influence of surface tensions on TRAAK conformational distribution. **a,** A schematic illustration depicting the down–up conformational transitions of the TRAAK channel under varying surface tensions. **b,** Representative dRMS trajectories of TRAAK relative to the down reference structure under a range of surface tensions (0, 5, 10, 20 and 30 mN/m). **c,** Free energy profiles of TRAAK under different surface tensions. Data from all trajectories, excluding the initial 1000 ns of each, are used for analysis. The shaded area surrounding each line represents the statistical uncertainty of the free energy, estimated using bootstrap analysis. **d,** Relationship between the fraction of the up state and surface tension. The dashed line represents a fitted curve obtained by applying a Boltzmann sigmoidal function to the simulation data.

We calculated the fraction of up states in the total simulations as a function of surface tension. The data aligned remarkably well with a Boltzmann sigmoidal function (Fig. 7d). From this curve, we identified the T50 (the surface tension at which 50% of the channels are in the up state) to be 8.2 mN/m. This value is somewhat higher than the experimental T50 of 4.4 mN/m, ^64^ which relates the current to the surface tension. It is important to note that a direct comparison between our simulated T50 and the experimental value is challenging for two reasons. Firstly, our simulations measure the fraction of up states, while experiments measure the ionic current. The relationship between TRAAK states and the current of the ion channel is complex. ^60–62^ Secondly, the structures used in our simulations are truncated, with parts of TM4 unresolved and therefore not included. This could result in a decreased interaction interface between TRAAK and the environment, potentially reducing the sensitivity to surface tension. Despite these limitations, our method clearly reveals the underling molecular mechanism of membrane surface tension-induced channel gating, which involves complex interactions between TRAAK and surrounding lipids under varying surface tensions.

## Discussion

Conformational transitions between different states are essential for proteins to exert their biological functions. Elucidating the detailed mechanisms by which proteins alter their structures to fulfill their roles remains a significant challenge in the field of structural biology and biophysics. Several key questions drive research in this field: (1) What are the pathways and mechanisms by which proteins transition between different conformational states? (2) How can we identify and quantitatively characterize important states along these transition pathways? (3) How do environmental factors, such as lipid composition, ligand binding, or changes in surface tension, modulate the conformational ensemble of proteins? All-atom MD simulations are one of the most widely-used computational methods to address these challenges. However, their application in this field is hindered by high computational costs and limited simulation timescales. ^9^ To overcome these limitations, in this study, we have developed the Multiple-basin Gō-Martini method. This approach capitalizes on the conformational flexibility of Gō-like models and the detailed environmental interactions, especially protein-lipid interactions, as defined in Martini force fields. Our method can simulate conformational transitions between two states of proteins and investigate the environment around proteins, particularly the biomembrane, in modulating the conformation distributions of proteins. Therefore, it overcomes two limitations of existing methods: the implicit environment representation in traditional Gō-like models and the single-basin restrictions in the Martini model.

To validate the accuracy and assess the capabilities of our method, we selected five diverse systems ranging from soluble to membrane proteins, from collective domain movements to fold-switching proteins, from natural to *de novo* designed proteins, and from standard environments to surface-tension systems. Our results demonstrate that the method can not only reliably simulate various protein conformational transition processes and capture the important intermediate states but also elucidate environmental modulation effects on protein state distributions. In addition, benchmark tests using the GlnBP system reveal that conformational transition rates in our method are 6–97 times faster than those in all-atom models. Considering the actual computational speed, the same system simulated with the Martini force field using our method achieves around 7,500 ns/day, whereas all-atom simulations using GROMACS on the same workstation yield about 250 ns/day. Consequently, the combination of these factors results in a speed-up of approximately 180 times using the Multiple-basin Gō-Martini method compared to those based on conventional all-atom simulations.

On the other hand, compared to implicit solvent Gō-like models, our method can provide insight into how environmental factors modulate protein state distributions, with a particular focus on protein-lipid interactions during protein conformational transitions. In this study, we investigate the surface tension-induced down–up transitions of TRAAK and for the first time present complete transition cycles under physiological surface tension (∼10 mN/m). Our findings reveal that lipid head groups, rather than tails, can insert into the fenestration formed by the TM4 and TM2 helices when TRAAK is in the down state. We also elucidate the detailed process of lipid evacuation from the fenestration as the protein transitions from the down to the up state. These results demonstrate the significant advantages of the Multiple-basin Gō-Martini method in simulating protein-lipid interactions during protein conformational transitions.

Although our method has successfully demonstrated its capability to elucidate the detailed conformational mechanisms of proteins, it is important to acknowledge its limitations. Firstly, the Multiple-basin Gō-Martini method requires structures of two distinct states of a given protein. This prerequisite limits its applicability to proteins with known multiple conformations, excluding those with only one known state or no structural information. This constraint restricts the broader application of the method to complex protein systems. However, this limitation could potentially be mitigated by incorporating deep learning methods to predict multiple protein states. ^68,69^ Secondly, our method needs a complex fine-tuning process to determine the optimal mixing parameters for coupling two energy basins and constructing the double-basin energy landscape. These parameters must be carefully adjusted to accurately reflect the thermal dynamics properties of protein systems. This time-consuming optimization process impedes the rapid application of our method to diverse systems. To address such challenges, Shinobu and co-workers have developed an accelerated parameter search method, which significantly expedites this process. ^37^ Thirdly, our current approach of handling side-chain flexibility introduces some limitations. To mitigate steric hindrance issues arising from restrained side chains during protein conformational changes, we temporarily remove the additional angle and dihedral restraints between side chains and backbones imposed by the -scFix flag in Martini models, which allows for enhanced flexibility and relaxation of side chains (see the Materials and Methods section for details). Although this approach allows for broader conformational sampling of side chains, it may impact the geometry of binding pockets and potentially affect the accuracy of protein-ligand binding simulations. However, with these limitations kept in mind, our method still provides highly valuable insight into protein conformational transitions and their environmental interactions.

The ongoing research aims to address several challenges, particularly focusing on expanding the accuracy and applicability of the Multiple-basin Gō-Martini method in future studies. Although this method provides valuable free energy profiles, it inevitably loses some intricate details due to the coarse-grained model. Thus, we propose combining the Multiple-basin Gō-Martini method with the coarse-grained to all-atom (CG2AT) resolution transformation ^70,71^ and enhanced sampling techniques. ^72–74^ This integration will allow us to re-sample along essential transition paths, revealing a more quantitative energy landscape of proteins. The second objective is to leverage this method to investigate the cooperative processes of multiple proteins. In previous Martini models, proteins are almost static and oscillate within local energy basins. Our method significantly expands the capabilities of Martini models by introducing spontaneous conformational transitions between different states, enabling a more dynamic exploration of how various proteins interact and collaborate to perform complex tasks within the cell. Additionally, while deep learning techniques have demonstrated remarkable progress in predicting various structural states of proteins with high accuracy, ^68,69,75–77^ and some works using deep learning-based methods to directly predict conformational distributions of proteins, ^78,79^ there is still substantial room for improvement. We believe that our method can improve the precision of these machine learning-based models by providing a wealth of reliable trajectory data sets that capture the thermodynamics of proteins, thus facilitating more accurate predictions of protein conformational space and a deeper understanding of the function mechanisms of dynamic proteins. ^80^

## Methods

### The Multiple-basin Gō-Martini Model

We use the Gō-Martini model ^31^ with the Martini 3 coarse-grained force field (version 3.0.0) ^25^ as the single-basin potential for different states of multiple-state proteins and use the macroscopic exponential mixing scheme^18^ to mix these states to construct the complex multiple-basin potential and to build the real energy landscape of proteins. The mixing scheme function is as follows:

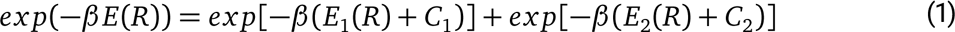

where *E*_1_ and *E*_2_ are the single-basin potentials to be mixed, *E* is the multiple-basin potential we need, *R* is the coordinates of one protein, and *β*, *C*_1_ and *C*_2_ are the mixing parameters which should be fine-tuned. *β* is related to the height of the energy barrier between the two basins, while *C*_1_ and *C*_2_ determine the energetic offsets between two basins. In this work, for the sake of simplicity, *C*_2_ is set to 0 throughout all protein systems and *C*_1_ is tuned regarding our requirements. Although this mixing scheme showcases a double-basin potential, this function can be extended to represent any number of basins, with *exp*[−*β* (*E_i_* (*R*) + *C_i_*)] added for the *i_th_* additional basin.

In the current work, we make the following combination rules in order to make the single-basin Gō-Martini models compatible for the exponential mixing scheme and reduce the calculation cost in MD simulations. The bond length is taken as the average length over the two reference states. If the bond type is the constraint in one state, while it is the harmonic bond in the other state, the final type is the harmonic bond. The backbone angle is taken as the average value over the two reference states if the angle difference between the two states is smaller than 15 degrees. If the angle type is the restricted angle in one state, while it is the g96 angle type in the other state, the final type is the restricted angle type. The backbone dihedral is taken as the average value over the two reference states if the difference between two states is smaller than 30 degrees. For contacts with a difference in sigma values less than 0.06 nm between two states, ^20^ the average value is used. If two interactions from two different states are combined, the parameters controlling the strength of the interactions are taken as the average values. Interactions that remain distinct between the two states after combination employ the above mixing scheme to construct the multiple-basin potential.

### General Simulation Settings

All simulations in our work were performed using the Martini 3 coarse-grained force field (version 3.0.0) ^25^ with the OpenMM MD package (version 8.1). ^81^ Details about generating protein CG models and setting up simulation systems are given in the following sections. For MD simulations, the systems were minimized and then equilibrated in the NPT ensemble with position restraints (1,000 kJ/mol/nm^2^) on the backbone of the CG proteins. Production simulations were conducted at a constant semi-isotropic or isotropic pressure of 1 atm and a temperature of 310 K depending on whether the systems contained the membrane or not. The Langevin integrator ^82^ with a friction coefficient of 1 ps^-1^ was used to maintain the temperature of the systems and the semi-isotropic or isotropic Monte Carlo membrane barostat ^83,84^ with a coupling frequency of 100 steps was used to maintain the pressure of the systems. The electrostatic interaction and the van der Waals interaction were both cut off at 1.1 nm and the long-range electrostatic interaction was calculated using the reaction field method ^85^ with a relative dielectric constant (epsilon_r = 15). All simulation parameters for running CG systems are listed in Supplementary Table S1. In addition, we have provided tutorials to show how to generate the Multiple-basin Gō-Martini model and run such MD simulations using OpenMM in our GitHub repository.

### Martini Models and System Setup

#### General Settings for Single-basin Gō-Martini Models

The Gō-Martini topology of each protein was generated using the tool martinize2 v0.96 to construct the single-basin potential. Notably, the -scFix flag in martinize2 was omitted to avoid generating additional angles and dihedrals between side chains and backbones, which could potentially distort the multiple-basin potential in our work. Additional Lennard-Jones interactions for the Gō-like model were added based on the Overlarp (OV) and replusive Contacts of Structural Units (rCSU) contacts (http://info.ifpan.edu.pl/~rcsu/rcsu/index.html). These interactions were set up with a distance cutoff of 0.3-1.1 nm and a dissociation energy of *ε* = 12.0 kJ/mol. The long-term external constraints applied to the secondary structure of the *β* -sheet were also converted into Lennard-Jones contact interactions with the same parameters, while the short-term constraints were excluded to avoid redundant interactions in the Gō-like model.

#### Glutamine Binding Protein

The open- and closed-state CG models of glutamine binding protein (GlnBP) were built using the Martini 3 force field, based on structures with PDB codes 1GGG ^41^ and 1WDN, ^40^ respectively. The sequences of both proteins were trimmed to have the same length (residues 5-224). Topologies of both states of GlnBP were generated following the process described in the previous section. The Multiple-basin Gō-Martini model of GlnBP was then constructed by combining the open- and closed-state single-basin Gō-Martini models, employing the aforementioned combination rules and the exponential mixing scheme. For our simulations, the open state of GlnBP was chosen as the initial conformation. The CG protein was placed in a rectangular box (8 × 8 × 7 nm^3^) filled with water using the program insane.py. ^86^ To mimic physiological conditions, 150 mM NaCl was added to keep the charge of the system neutral. To determine the optimal parameter setting, a series of values of parameter *β* (1/100, 1/200, 1/300, 1/400, and 1/500 mol/kJ) and parameter *C*_1_ (-100, -200, -300, -400, and -500 kJ/mol) were tested, resulting in a total of 25 distinct parameter combinations. For each combination, a 5-µs simulation was conducted. Based on these trail tests, the optimal parameters (*β* =1/300 mol/kJ, *C*_1_=-300 kJ/mol, *C*_2_=0 kJ/mol) were identified. Using this set of parameters, 10 independent CG simulations were performed, each running for 20 µs, to investigate the spontaneous conformational transitions of GlnBP. Additionally, to assess the influence of the initial state and ensure the convergence of our simulations, another series of simulations with the same settings but with the closed state as the initial state were also performed.

#### Arc

The sheet and helix state CG models of Arc were built using the Martini 3 force field, based on structures with PDB codes 1ARR ^46^ and 1QTG, ^47^ respectively. Because the Arc protein with the N11L mutation has been reported to exist in two states simultaneously, both 1ARR (wild-type) and 1QTG (including the N11L and L12N mutations) were modified to include only the N11L mutation using Modeller. ^87^ The sequences of both proteins were trimmed to have the same length (residues 1-53 for each chain). Topologies of the sheet and helix states of Arc were generated following the process described in the previous section. The Multiple-basin Gō-Martini model of Arc was then constructed by combining the sheet- and helix-state single-basin Gō-Martini models, employing the aforementioned combination rules and the exponential mixing scheme. For our simulations, the helix state of Arc was chosen as the initial conformation. The CG protein was placed in a cubic box (6.5 × 6.5 × 6.5 nm^3^) filled with water using the program insane.py. ^86^ To mimic physiological conditions, 150 mM NaCl was added to keep the charge of the system neutral. After parameter optimization, the suitable parameters (*β* =1/350 mol/kJ, *C*_1_=25 kJ/mol, *C*_2_=0 kJ/mol) were determined. With this set of parameters, 10 independent CG simulations were performed, each running for 20 µs, to investigate the spontaneous conformational transitions of Arc.

### *De novo*-designed Hinge

The bound and unbound state CG models of Hinge (cs207) were built using the Martini 3 force field, based on structures with PDB codes 8FIQ and 8FIN, ^50^ respectively. The sequences of both proteins were trimmed to have the same length (residues 2-173). Topologies of the bound and unbound states of Hinge were generated following the process described in the previous section. The Multiple-basin Gō-Martini model of Hinge was then constructed by combining the bound- and unbound-state single-basin Gō-Martini models, employing the aforementioned combination rules and the exponential mixing scheme. For our simulations, the bound state of Hinge was chosen as the initial conformation. The substrate peptide in the binding pocket of Hinge was converted to the Gō-Martini model, following the same protocol used for Hinge, with the addition of the -scFix flag in martinize2 to maintain the stability of the side chains. The contact interactions between Hinge and the peptide were defined based on the OV and rCSU contact maps, with a distance cutoff of 0.3-1.1 nm and a dissociation energy of *ε* = 12.0 kJ/mol. The CG complex was placed in a cubic box (8 × 8 × 8 nm^3^) filled with water using the program insane.py. ^86^ To mimic physiological conditions, 150 mM NaCl was added to keep the charge of the system neutral. After parameter optimization, the suitable parameters (*β* =1/500 mol/kJ, *C*_1_=-520 kJ/mol, *C*_2_=0 kJ/mol) were determined. With this set of parameters, we then performed 10 independent CG simulations. To investigate the unbinding process, each trajectory consisted of 10 µs standard MD, followed by 500 ns of steered MD, and another 9.5 µs of standard MD. During the pulling step, we applied a constant-velocity force (0.008 nm/ns and 1000 kJ/mol/nm²) between the centers of mass of the backbone atoms of the peptide and the Hinge binding pocket (residues 67-132).

To study the binding process, a second system using the unbound state of Hinge as the initial conformation was constructed, with 10 peptides randomly distributed in the simulation box (8 × 8 × 8 nm^3^). After adding 150 mM NaCl, we performed 10 independent CG simulations, each running for 20 µs using identical multiple-basin mixing parameters.

#### SemiSWEET

The outward-open and inward-open state CG models of SemiSWEET were built using the Martini 3 force field, based on structures with PDB codes 4X5N and 4X5M, ^53^ respectively. The sequences of both proteins were trimmed to have the same length (residues 2-91 for each chain). Topologies of the outward-open and inward-open states of SemiSWEET were generated following the process described in the previous section. The Multiple-basin Gō-Martini model of SemiSWEET was then constructed by combining the outward-open and inward-open states of single-basin Gō-Martini models, employing the aforementioned combination rules and the exponential mixing scheme. For our simulations, the outward-open state of SemiSWEET was chosen as the initial conformation. The CG protein was embedded into a complex bilayer within a rectangular box (7 × 7 × 8 nm^3^) filled with water using the program insane.py, ^86^ in which the membrane contained 80% POPE and 20% POPG. To mimic physiological conditions, 150 mM NaCl was added to keep the charge of the system neutral. After parameter optimization, the suitable parameters (*β* =1/1000 mol/kJ, *C*_1_=-840 kJ/mol, *C*_2_=0 kJ/mol) were determined. With this set of parameters, 10 independent CG simulations were performed, each running for 20 µs, to investigate the spontaneous conformational transitions of SemiSWEET.

#### TRAAK

The down- and up-state CG models of TRAAK were built using the Martini 3 force field, based on structures with PDB codes 7LJB^65^ and 4WFE, ^57^ respectively. The mutations and missing loops of 7LJB (mutations G158D, N104Q, N108Q, and residues 104-109, 285-286) and 4WFE (mutations N104Q, N108Q, and residues 103-109) were filled using Modeller. ^87^ The sequences of both proteins were trimmed to have the same length (residues 28-286 for chain A and B). Topologies of the down and up states of TRAAK were generated following the process described in the previous section. The Multiple-basin Gō-Martini model of TRAAK was then constructed by combining the down and up states of single-basin Gō-Martini models, employing the aforementioned combination rules and the exponential mixing scheme. For our simulations, the down state of TRAAK was chosen as the initial conformation. The CG protein was embedded into POPC bilayer (146 and 152 POPC molecules in the upper and lower leaflets) within a rectangular box (11 × 11 × 12 nm^3^) filled with water using the program insane.py. ^86^ To mimic physiological conditions, 150 mM NaCl was added to keep the charge of the system neutral. After parameter optimization, the suitable parameters (*β* =1/215 mol/kJ, *C*_1_= -30 kJ/mol, *C*_2_=0 kJ/mol) were determined. To investigate the influence of surface tension on the state distribution of TRAAK, a series of surface tension values of 0, 5, 10, 20, 30 mN/m were applied to the membrane. For each condition, 10 independent CG simulations were performed, each running for 20 µs.

### Atomistic Simulations

For our study, we selected GlnBP as the reference protein to compare the results between allatom and coarse-grained simulations. The prepared structures of open and closed states of GlnBP used in the CG simulations were submitted to the CHARMM-GUI server ^88^ to generate allatom systems. The proteins were placed in the center of the simulation boxes and solvated with TIP3P water molecules. To neutralize the system’s charges, 150 mM NaCl was added to each box. All simulations were performed using GROMACS 2021.2^89^ with the CHARMM36m force field. ^90^ The systems were first energy minimized using the steepest-descent algorithm for 5000 steps. They were then equilibrated for 0.125 ns in the NVT ensemble and 1 ns in the NPT ensemble, with a restraint of 400 and 40 kJ/mol/nm² applied to the backbone and side chain atoms, respectively. During the NVT equilibration stage, all systems were coupled to the Berendsen thermostat ^91^ to maintain the simulation temperature at 310 K. In the NPT equilibration, the Berendsen barostat was added for isotropic pressure coupling to set the pressure at 1 bar. Three independent 1000 ns production simulations were conducted for each state. During the production stage, the V-rescale thermostat ^92^ and the Parrinello-Rahman barostat ^93^ were used to maintain the temperature at 310 K and the pressure at 1 bar, respectively.

### Simulation Analysis

The distance root-mean-square displacement (dRMS) was calculated using the following equation:

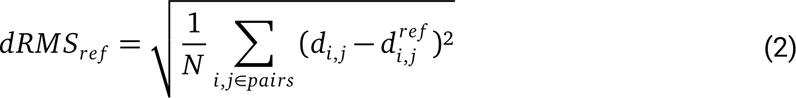

where *d_i_*,*_j_* and 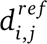 are the distance between backbone atoms of residues i and j in the protein and the reference state, respectively. N is the number of essential residue pairs in the reference structures that are separated by at lease four residues in the sequence and 6–50 Å in space, and exhibit a distance difference of at least 5 Å between the two reference states in the Multiple-basin Gō-Martini models. ^37^

The one-dimensional (1D) and two-dimensional (2D) free energy surfaces in our simulations were defined using the following equation:

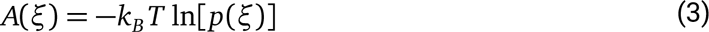

where *A*(*ξ*) is the free energy as a function of the reaction coordinate *ξ*, *k_B_* is the Boltzmann constant, *T* is the simulation temperature, and *p*(*ξ*) is the probability distribution of states along *ξ*. In our analysis, the first 1000 ns of each trajectory was discarded to avoid the influence from the initial state of the simulations. To facilitate comparison, we subtracted the minimum value of the free energy from the entire profile, setting the global minimum to zero. The statistical uncertainty of the free energy profiles was estimated using bootstrap analysis with 200 times of resampling the raw data. The minimum free energy path of the 2D free energy surfaces was obtained using the MEPSAnd software. ^94^

The equilibrium constant *K_eq_* between two states was defined as:

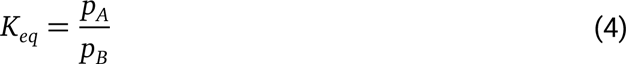

where *p_A_* and *p_B_* represent the populations of frames in state A and state B, respectively. The MD trajectory frames were assigned to their respective states on the basis of the reaction coordinate values, with the cutoff values at the location of the barriers in the free energy profile.

Principal component analysis (PCA) was calculated with MDAnalysis. ^95^ To follow the principles of all-atom to coarse-grained mapping, we only considered the center of mass of backbones in all-atom models. Trajectories from all-atom simulations of both open-state and closed-state GlnBP were analyzed to identify the large-amplitude conformational changes. To evaluate the overlapping consistency between all-atom and coarse-grained simulations, a representative trajectory of the CG simulations was projected onto the first two principal components (PCs) derived from the all-atom simulations.

Additionally, distance calculation, protein-lipid interactions and RMSD were analyzed with MDAnalysis. ^95^ All structural figures and movies were generated using ChimeraX. ^96^

## Data Availability

The codes and tutorials for the Multiple-basin Gō-Martini simulations can be found on Github for public access after peer review.

## Acknowledgements

We thank Dr. Yuji Sugita at RIKEN for insightful suggestions.

## Author Contributions

S.Y. and C.S. conceived the idea and designed the research. S.Y. conducted simulations and analyzed data. S.Y. and C.S. wrote the manuscript. C.S. acquired funding and supervised the work.

## Supplementary Information

### 1 Supplementary Tables

**Table S1:**
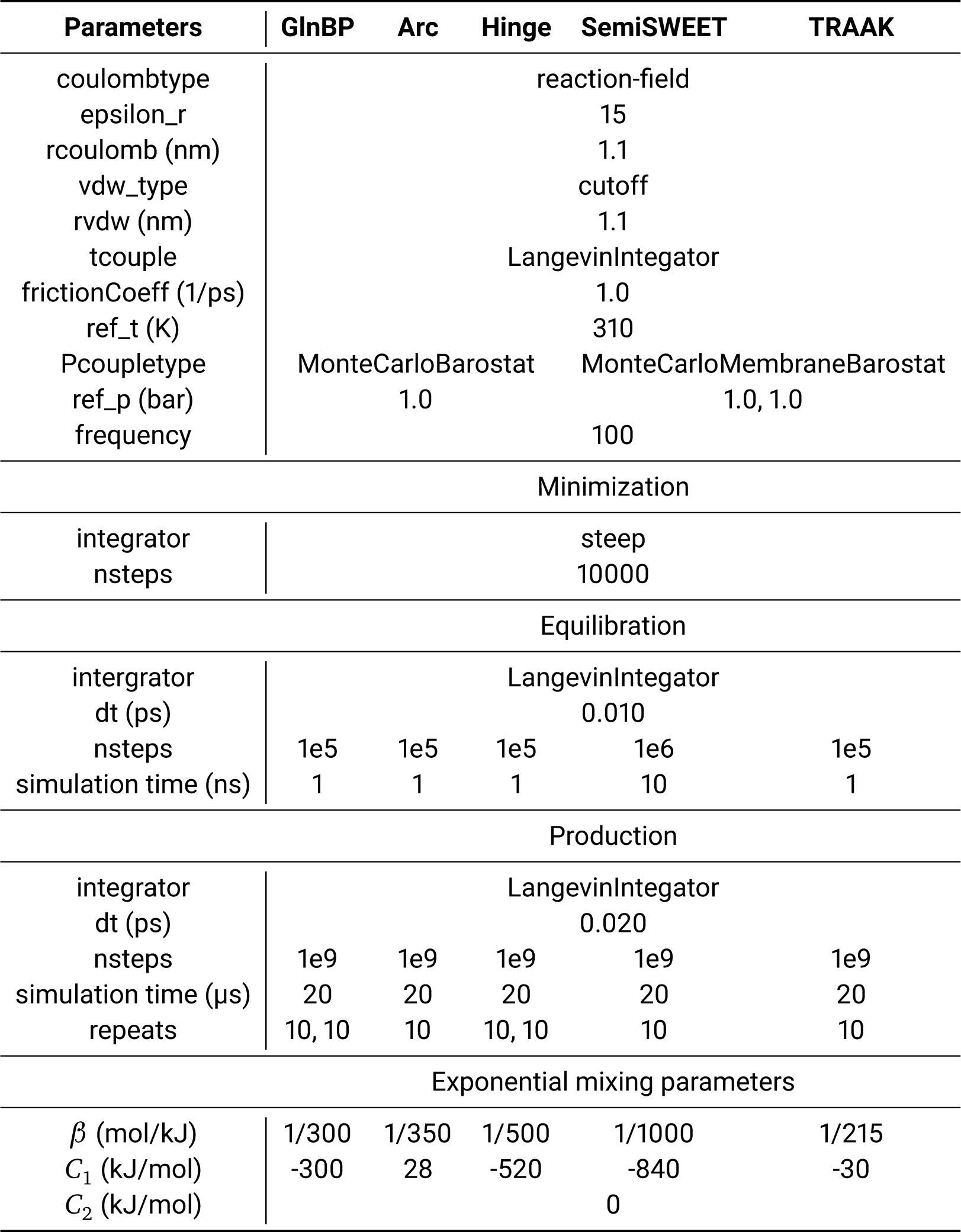
Parameters and summary for Multiple-basin Gō-Martini simulations in this work.

### 2 Supplementary Figures

**Figure S1:**
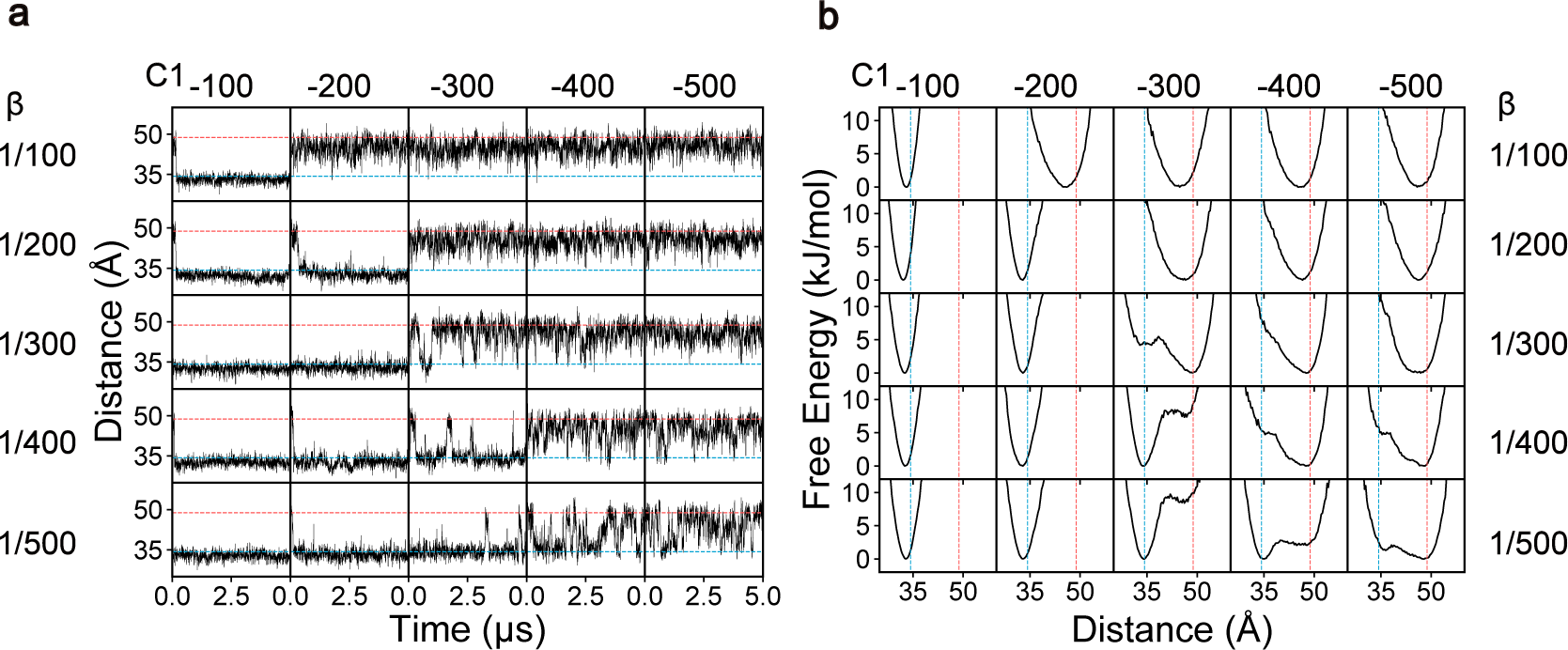
Coupling parameter search for GlnBP conformational transitions. **a,** Time evolution of the distance between backbone atoms of T59 and T130 under different coupling parameters. **b,** Free energy profiles of GlnBP conformational changes under corresponding coupling parameters. The distance between T59 and T130 is used as the reaction coordinate, with each free energy curve generated from 5 µs of coarse-grained simulation data.

**Figure S2:**
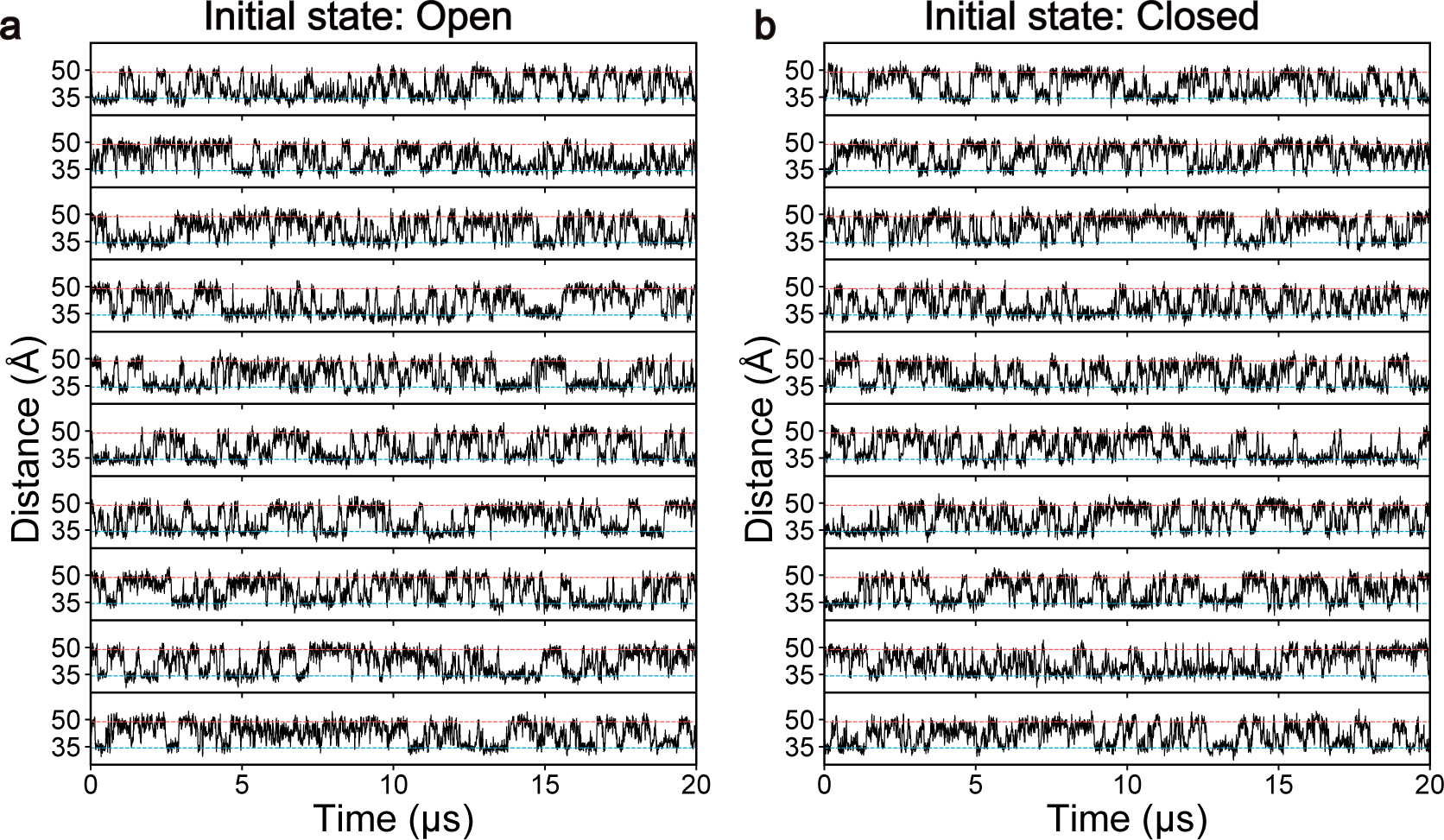
Multiple-basin Gō-Martini simulations for GlnBP. **a, b,** Time evolution of the distance between backbone atoms of T59 and T130 over the course of 10 independent simulations, initiated from open and closed states, respectively

**Figure S3:**
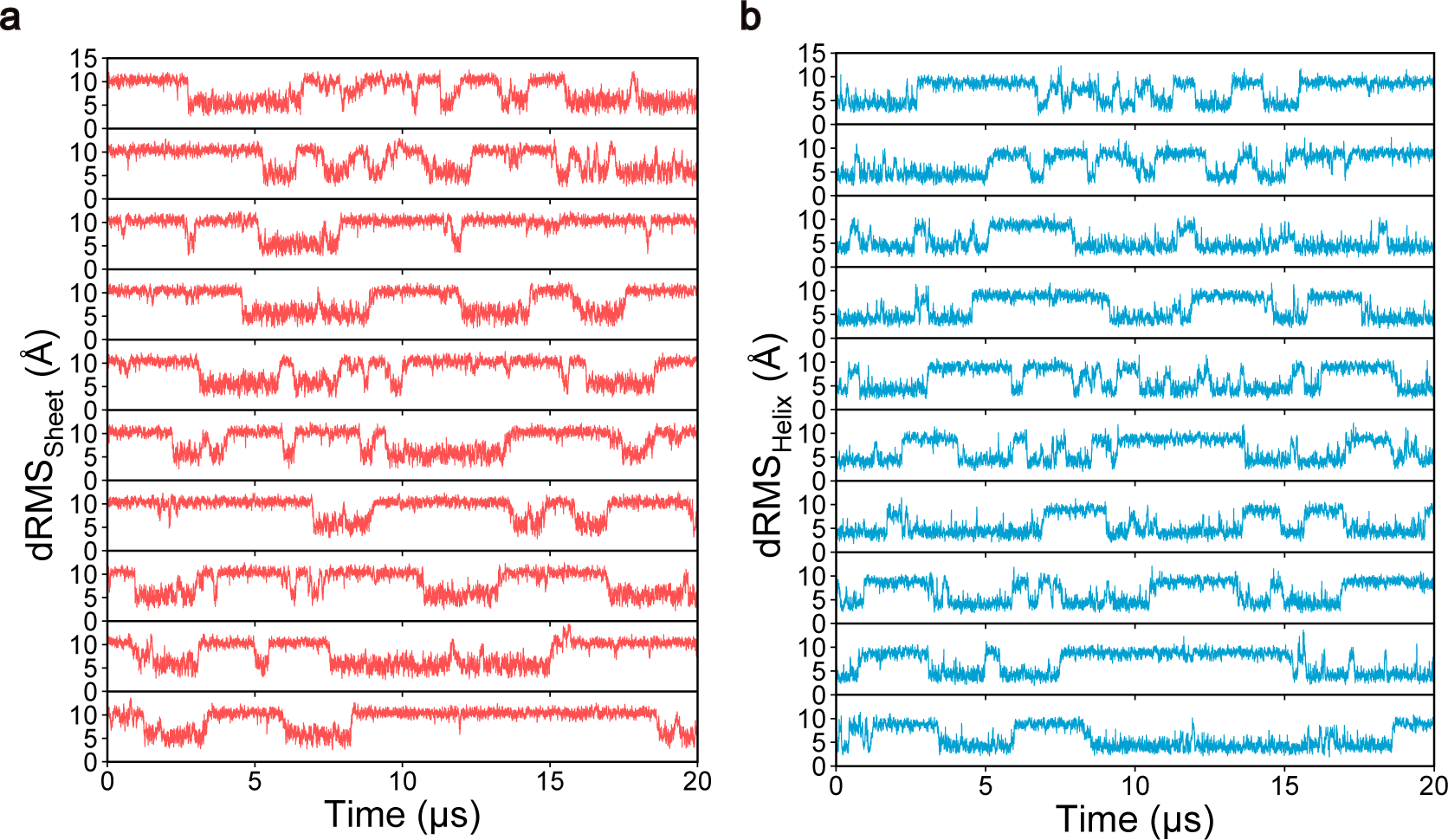
Multiple-basin Gō-Martini simulations for Arc. **a, b,** Time evolution of dRMS trajectories of Arc over the course of 10 independent simulations, relative to the sheet (red) and helix (blue) reference structures, respectively.

**Figure S4:**
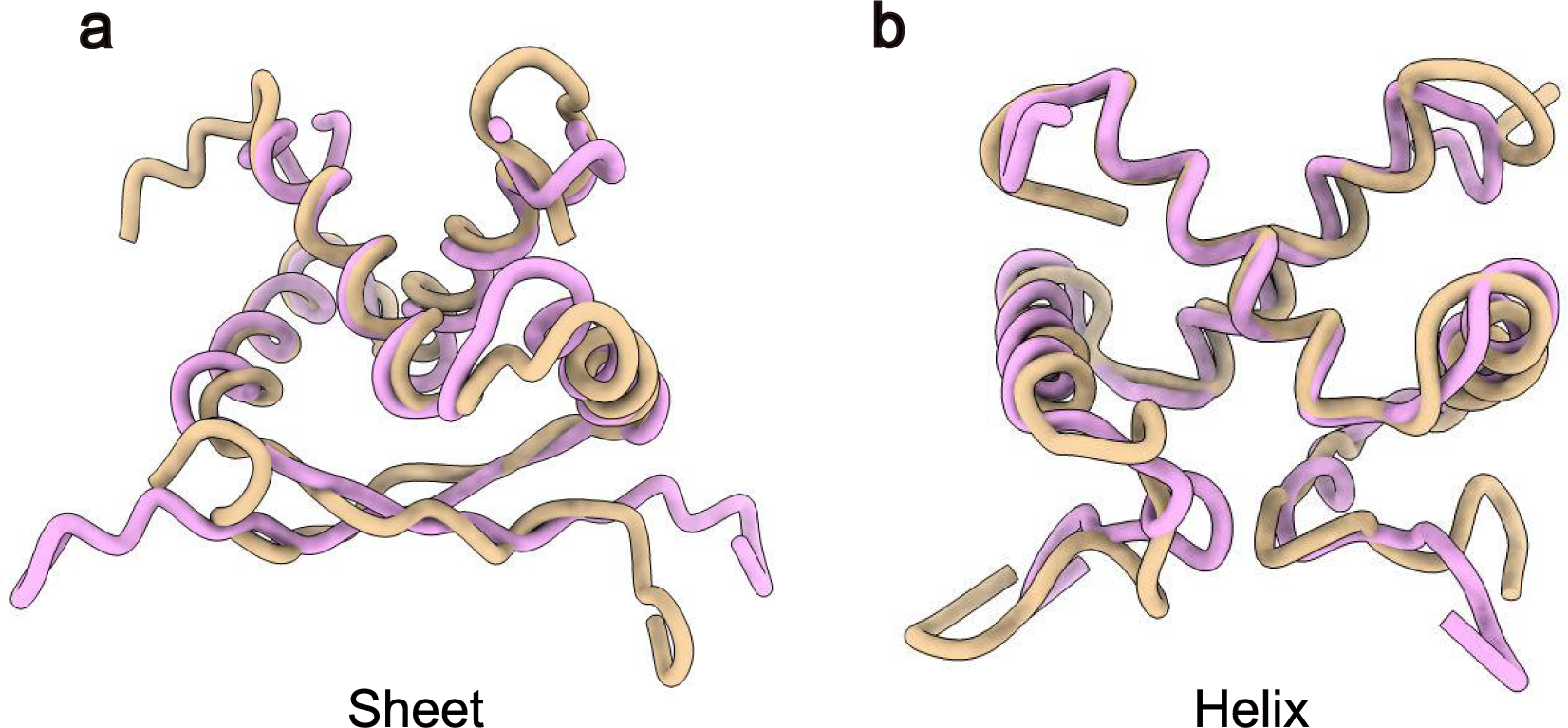
Representative basin structures of Arc. **a, b,** Representative structures of Arc in the helix and sheet basin states, respectively. Basin structures are depicted in tan, while reference structures are shown in plum.

**Figure S5:**
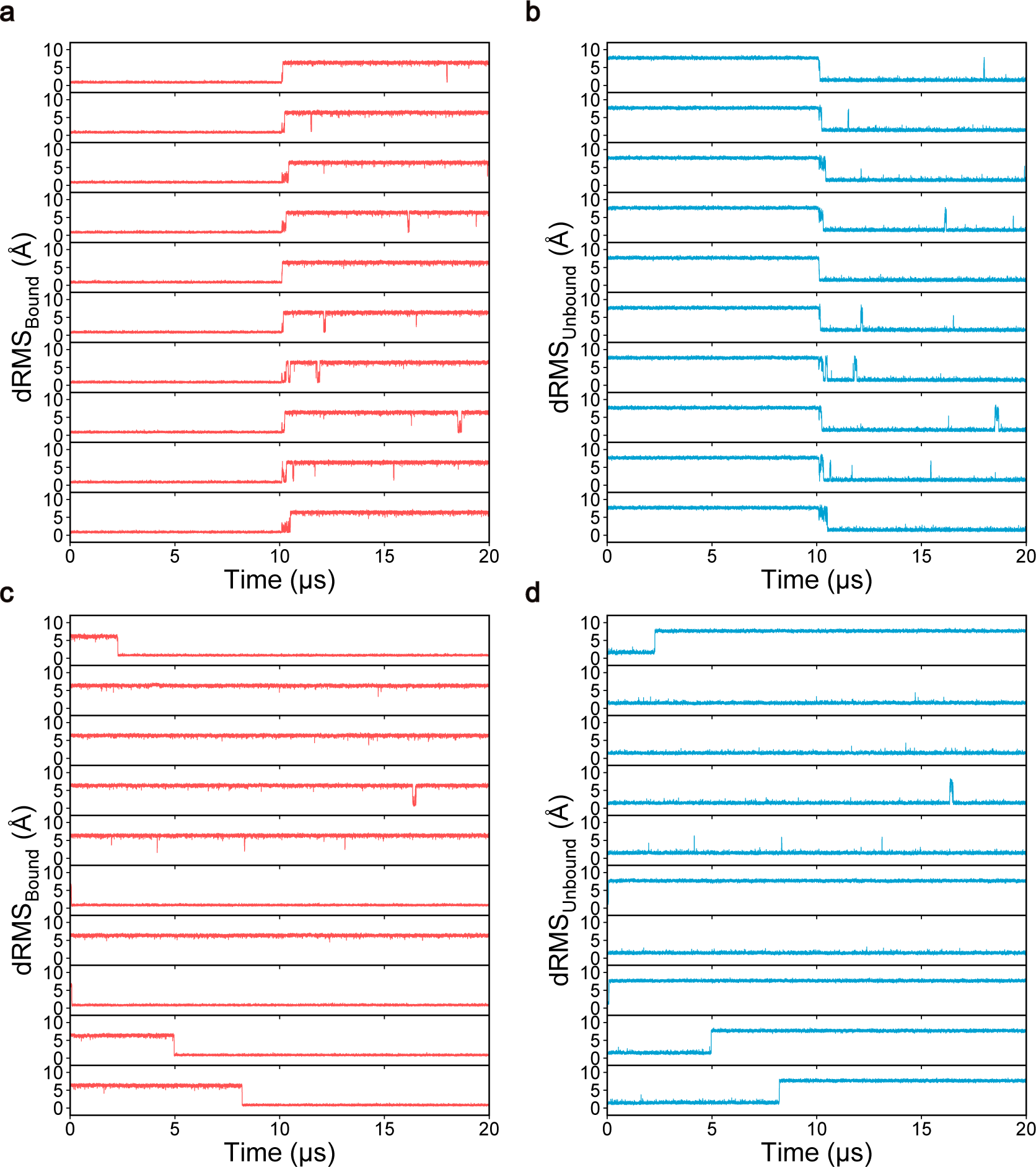
Multiple-basin Gō-Martini simulations for Hinge. **a, b,** Time evolution of dRMS trajectories of Hinge over the course of 10 independent unbinding simulations, relative to the bound (red) and unbound (blue) reference structures, respectively. **c, d,** Time evolution of dRMS trajectories of Hinge over the course of 10 independent binding simulations, relative to the bound (red) and unbound (blue) reference structures, respectively.

**Figure S6:**
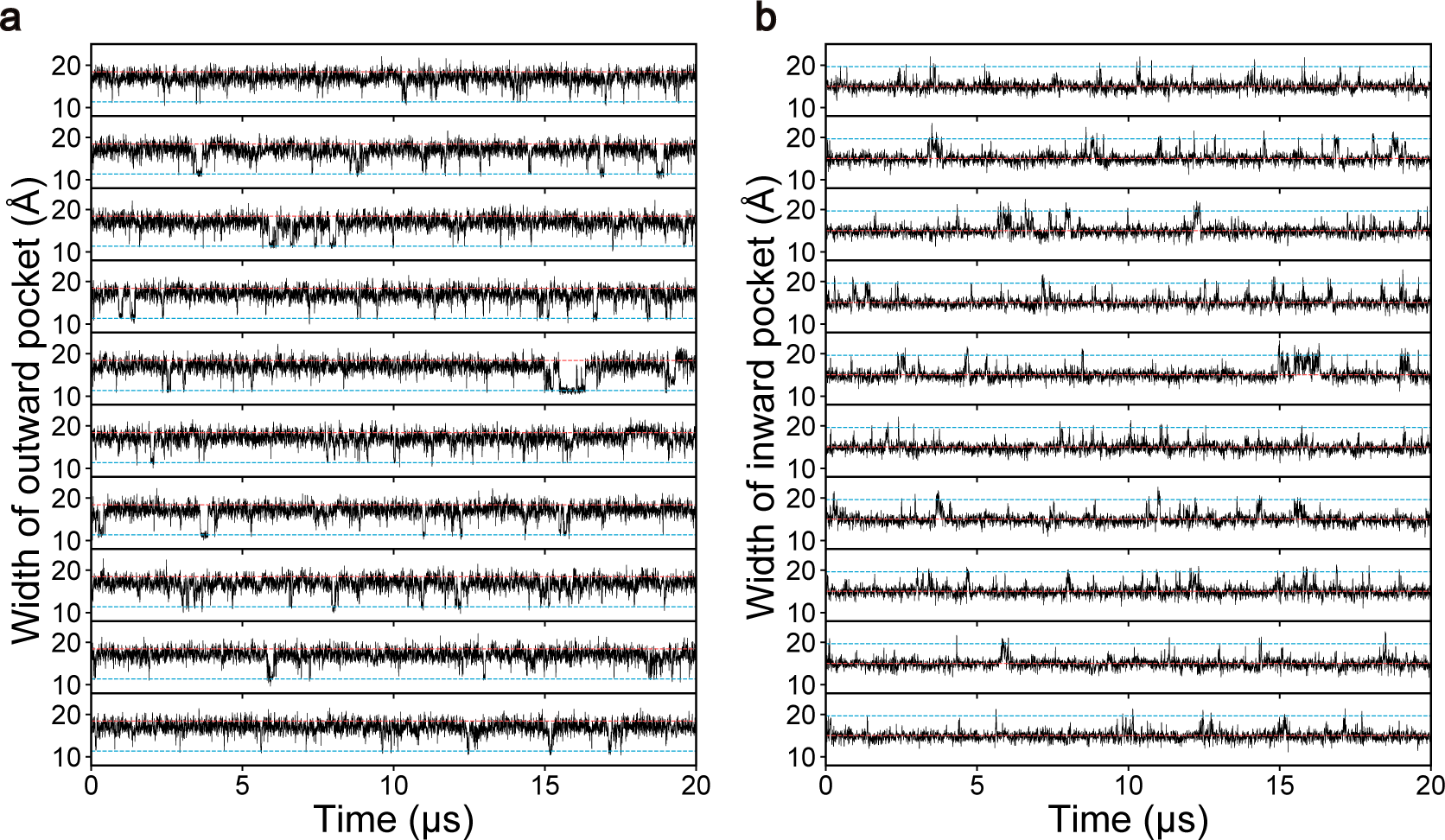
Multiple-basin Gō-Martini simulations for SemiSWEET. **a, b,** Time evolution of width of outward and inward binding pockets of SemiSWEET over the course of 10 independent simulations, respectively.

**Figure S7:**
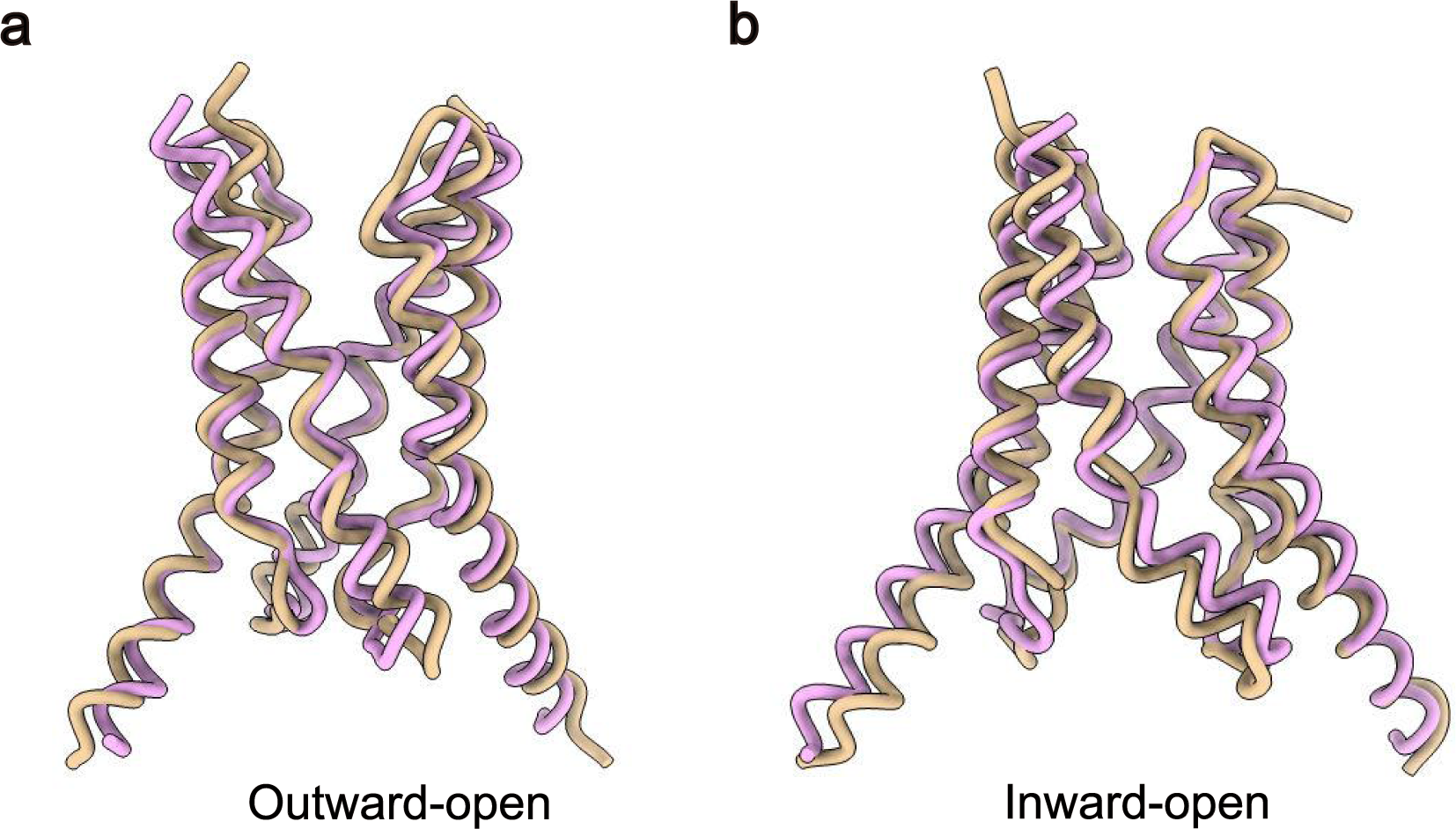
Representative basin structures of SemiSWEET. **a, b,** Representative structures of SemiSWEET in the outward-open and inward-open basin states, respectively. Basin structures are depicted in tan, while reference structures are shown in plum.

**Figure S8:**
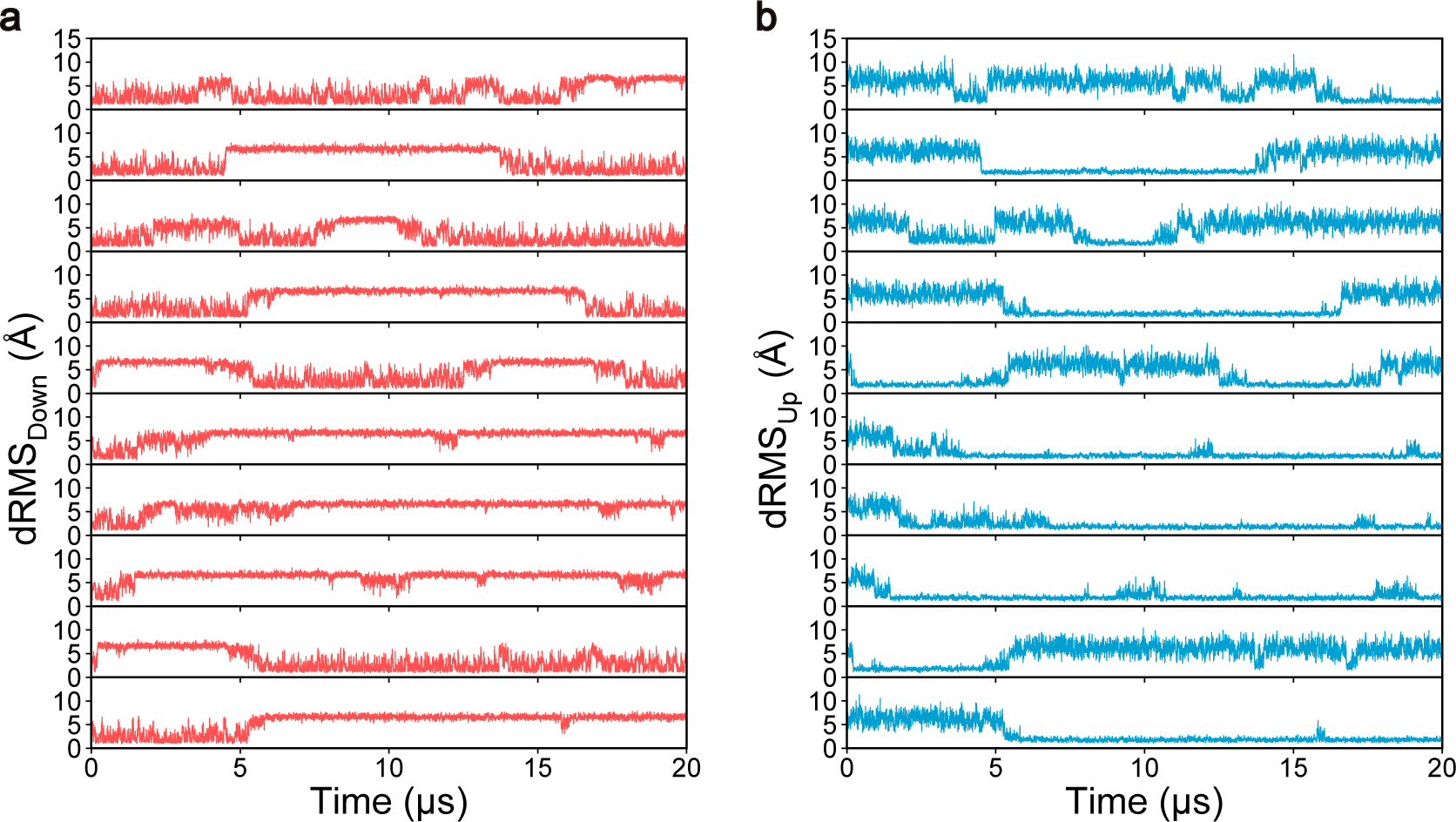
Multiple-basin Gō-Martini simulations for TRAAK. **a, b,** Time evolution of dRMS trajectories of TRAAK with a surface tension of 10 mN/m over the course of 10 independent simulations, relative to the down (red) and up (blue) reference structures, respectively.

**Figure S9:**
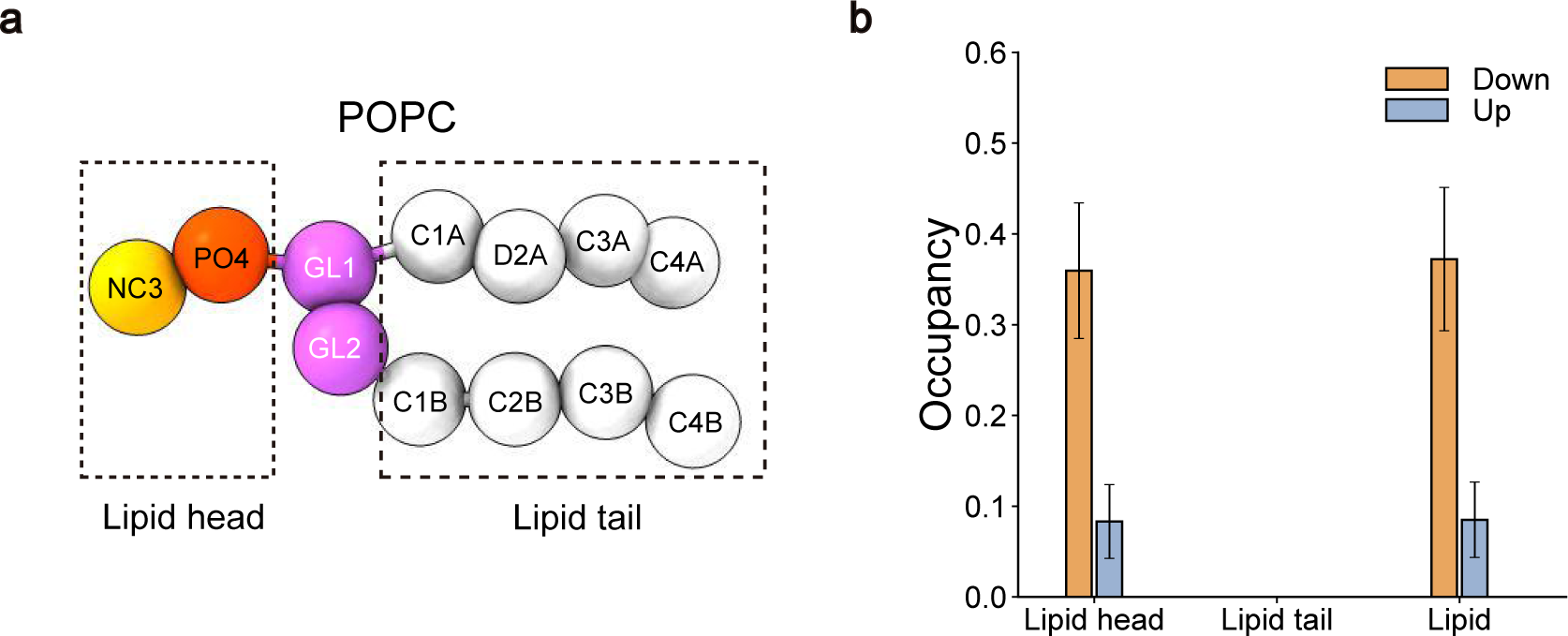
Lipid occupancy analysis of TRAAK. **a,** Coarse-grained POPC molecule in a ball-and-stick representation with dashed rectangles highlighting the lipid head and tail used in the analysis. **b,** Occupancy of lipid head group, lipid tail group and the entire lipid molecule in the down and up states of TRAAK. The occupancy is measured within an 8 Å radius from the center of the bottom region of the selective filter (T129 and T238 of chain A and B) in TRAAK. This analysis encompasses all 10 simulations, with error bars representing the standard deviation.

### 3 Supplementary Movie

**Supplementary Movie 1. Conformational transitions of GlnBP**. The movie shows a representative trajectory of open–closed transitions of GlnBP. The protein is shown as a ribbon, with the large domain colored sky blue and the small domain salmon. The remaining part is colored tan. The backbone atoms of T59 and T130 are highlighted with light green spheres.

**Supplementary Movie 2. Conformational transitions of Arc**. The movie shows a representative trajectory of sheet–helix transitions of Arc. The protein is shown as a tan ribbon.

**Supplementary Movie 3. Conformational transitions of Hinge during the peptide unbinding process**. The movie shows a representative trajectory of the bound-to-unbound transition of Hinge during the peptide unbinding process. The protein backbone of Hinge is shown in tan, with Helices 4-6 highlighted in sky blue. The substrate peptide is depicted as a salmon ribbon.

**Supplementary Movie 4. Conformational transitions of Hinge during the peptide binding process**. The movie shows a representative trajectory of the unbound-to-bound transition of Hinge during the peptide binding process. The protein backbone of Hinge is shown in tan, with Helices 4-6 highlighted in sky blue. The binding peptide is depicted as a salmon ribbon, while the other nine peptides are displayed in light green.

**Supplementary Movie 5. Conformational transitions of SemiSWEET**. The movie shows a representative trajectory of SemiSWEET transitioning between outward-open and inward-open states. The protein is shown as a tan ribbon, with the backbone atoms of M39 and D59 highlighted with salmon and light green spheres, respectively.

**Supplementary Movie 6. Conformational transitions of TRAAK**. The movie shows a representative trajectory of down–up transitions of TRAAK. The protein is shown as a tan ribbon, with the backbone atoms of P155 and I279 highlighted with light green and salmon spheres, respectively. Lipids binding to the fenestration formed by TM2 and TM4 are depicted using a ball-and-stick model, with the head groups colored orange and red while the remaining part colored middle orchid.

